# In-vitro digestion models to examine the effects of Plant-Based Meat substitutes on Gut Microbial Metabolites

**DOI:** 10.1101/2023.10.24.562654

**Authors:** David Izquierdo-Sandoval, Xiang Duan, Christos Fryganas, Tania Portolés, Juan Vicente Sancho, Josep Rubert

## Abstract

The increasing popularity of plant-based meat alternatives (PBMAs) has triggered a contentious debate about their impact on gut health in comparison to traditional animal-based meats. This study investigates the digestibility and bioavailability of a beef patty, a commercial PBMA, and a homemade pea protein-based ’patty’ by examining their influence on gut microbial metabolism. Fecal samples from five different donors were utilized to replicate colonic fermentation in vitro, with samples collected at various time points (0, 6, 12, 24, 32, and 48 hours). A rapid biochemical profiling, comparing red meat and meat analogs in terms of traditional biomarkers of gut health (ammonia, phenols, indoles, pH, and short-chain fatty acids), was conducted. Additionally, an untargeted metabolomics workflow specially designed for time-series studies, utilizing ultra-high performance liquid chromatography hyphenated to a quadrupole time-of-flight mass spectrometer (UPLC-QTOF MS), was implemented to assess differences in terms of protein-related gut microbial metabolites (GMMs). The findings of this approach revealed notable differences in the production of intestinal inflammation markers, metabolites related to the carnitine pathways, and GMMs with signaling functions in the intestinal tract during the fermentation of animal- and plant-based burgers.

## 1. INTRODUCTION

The consumption of animal-derived protein is dramatically increasing in emerging countries while barely decreasing in developed countries ^1^. This will bring us a scenario where meat consumption worldwide will impact greenhouse gas emissions and the demand for environmental resources ^2^. On top of that, excessive red meat intake has been associated with gastrointestinal (G.I.) diseases, including inflammatory bowel disease (IBD), colorectal cancer (CRC) ^3^, and non-communicable diseases (NCDs) such as diabetes type 2 and coronary heart diseases ^4,5^. As an alternative, plant-based meat analogs (PBMA) are gaining popularity as meat substitutes among health and environmentally-conscious consumers with similar organoleptic characteristics to traditional meat products. They are constituted of pure (or partially purified) plant proteins (e.g., soybean, pea, grain, etc.), vegetable oils, binding agents, and other ingredients such as preservatives and sweeteners ^6^. After being ingested and digested, both animal and plant proteins go through the G.I. tract, and in the upper and lower intestines, they are utilized by trillions of microorganisms that encode the ability to generate thousands of molecules ^7^. Bacterial proteases hydrolyze proteins and large peptides into small peptides and single amino acids (A.A.s), which in turn are fermented to produce short-chain fatty acids (SCFA), branched-chain fatty acids (BCFA), ammonia, amines, hydrogen sulfide, phenols, indoles, among other GMMs. Furthermore, the intestinal microbiota can also recycle nitrogen and synthesize A.A.s *de novo* ^8^. These microbial-related metabolites, either as a single agent or combinations, can positively or negatively affect intestinal epithelial cells and human health ^9^. GMMs are essential for the metabolic, immune, and neurological systems of the host ^10^. However, the impact of replacing animal-derived proteins with plant-based proteins via PBMA on GMM production remains unclear.

A recent interventional study, in which fecalsamples from participants fed with different animal protein sources (red meat, fish, poultry, eggs, and cheese) and pea-based meat were subjected to 16S rRNA sequencing, the authors reported an increase in butyrate producers associated with pea-based meat that may be potentially connected in the 4-aminobutyrate/succinate and glutamate metabolic pathways ^11^. Butyrate is a crucial SCFA since it is the primary energy source for colonocytes and activates intestinal glucogenesis, which positively impacts the host’s glucose and energy balance ^12^. Recently, Zhou and co-authors claimed that SCFA production was boosted in the ascending and descending colon when pea based PBMA was employed to feed the Simulated Human Intestinal Microbial Ecosystem (SHIME^®^) ^13^. Indeed, numerous studies have shown beneficial anti-inflammatory effects with soy and pea-based protein interventions ^14^.

In addition to SCFAs, an extensive repertoire of GMMs can be produced by consuming plant and animal proteins. Therefore, advanced analytical tools are highly required to solve this puzzle. Due to the chemical structure of many GMMs, ultra-high performance liquid chromatography coupled with high-resolution mass spectrometry (UHPLC-HRMS) has emerged as a fast, sensitive, and accurate technology for the screening of a virtually unlimited number of GMMs in a single analysis ^15^. The use of hybrid HRMS mass analyzers, such as quadrupole-time of flight (QTOF), enables the acquisition of data on ionized molecules and fragmented ions in a single injection, and therefore significantly increases the possibilities for metabolite/molecule/analyte identification ^16^. In our research work, an untargeted approach aimed to compare fingerprints of metabolites that were altered in response to different protein sources during an *in vitro* colonic fermentation. During this process, a massive volume of data per sample (thousands of features) is generated, making it paramount to select the right workflow and software tools to perform efficient and rigorous metabolomic data analysis, including data processing steps (peak picking, deconvolution, peak matching, and peak alignment across the samples), annotation, and statistical analysis ^17–20^.

This work used a beef patty, a commercial PBMA (pea-based meat), and a homemade pea protein-based ’patty’ to investigate GMM profiles. These food matrices were first cooked and digested using the INFOGEST protocol. The undigested fractions were the starting materials to generate fecal batch cultures. Fecal materials from five different donors were used to mirror colonic fermentation *in vitro*, collecting samples at different time points (0, 6, 12, 24, and 48 h). Protein-related GMMs were assessed through an untargeted metabolomics approach by using UHPLC-QTOF MS. Two different chromatographic columns were used, reversed-phase (R.P.) and hydrophilic interaction liquid chromatography (HILIC). The pipeline to process the generated data included several open software packages for data processing, retention time alignment, identification, and statistical analysis. Ammonia, pH, and total indole and phenol contents were assessed using different target methodologies.

## 2. MATERIALS AND METHODS

### 2.1. Chemicals and reagents

Porcine pepsin (P6887), porcine pancreatin (P1750, 4× USP), and porcine bile salt preparation (B8631) were purchased from Sigma-Aldrich (Merck KGaA, Germany). KCl, KH_2_PO_4_, MgCl_2_·(H_2_O)_6_, CaCl_2_·(H_2_O)_2_, and pure ethanol were purchased from VWR International B.V. (Netherlands). Yeast extract, peptone, mucine and L-cysteine HCl, NaCl, (NH_4_)_2_CO_3_, NaHCO_3_, NaOH, HCl, and Tween 80 were purchased from Sigma-Aldrich Chemie B.V. (Netherlands).

For the preparation of mobile phases for chromatographic analysis, LC-MS grade methanol (MeOH), acetonitrile (ACN), 2-propanol (IPA) and water were purchased from Sigma-Biosolve B.V. (Valkenswaard, Netherlands). For the rest of the experimental part Milli-Q water was produced by Milli-Q PURELAB Ultra, ELGA LabWater (Lane End, U.K.). Ammonium acetate, ammonium formate, formic acid, and methyl tert-Butyl ether (MTBE) were purchased from Sigma-Aldrich Chemie NV (Zwijndrecht, Netherlands). Internal standards (I.S.) were also purchased from Sigma-Aldrich Chemie NV (Zwijndrecht, Netherlands) and were added to the mixtures in the following concentrations: :500 ng mL^-1^ of tryptophand_5_, trans cinnamic acid d_7_, L-Lysine d_4_ hydrochloride, 250 ng mL^-1^ of dopamine d_4_, and ^13^C Glycocholic acid-(glycyl-1-^13^C), and 150 ng mL^-1^ of acrylamide-d_3_.

### 2.2. Burger material

Beef burger (125 g) was purchased from Slagerij Elings B.V. in Wageningen, consisting of 80% lean beef meat and 20% beef fat, this burger is labelled as RM. An available plant-based commercial burger, labelled as PBCB, was obtained from Albert Heijn. For the homemade plant-based burger, labelled as P.P., pea protein isolate was provided by Ingredion (Hamburg, Germany), and coconut oil was purchased from Thermo Fisher Scientific (Breda, Netherlands). The protein, fat, and water ratios were similar in the three products. More details on the composition of the foodstuff material can be bound in Table S1. The three burger types were baked in a conventional oven for 6-10 minutes at 180°C until the core reached 60°C. (More details in the Supporting Information, Section 1.1).

### 2.3. Simulated *in vitro* gastrointestinal digestion

The three baked patties were digested using the INFOGEST method ^21,22^ consisting of a simulated oral, gastric, and intestinal phase, with modifications. The compositions (%, w/w) of the simulated salivary fluid (SSF), simulated gastric fluid (SGF, pH 3.0 ± 0.05), and simulated intestinal fluid (SIF, pH 7.0 ± 0.05) were as reported in previous works ^23^. The entire process was carried out at a constant temperature of 37 °C and with 5 replicates per patty (More details in the Supporting Information, Section 1.2). At the end of the process, samples were incubated for 2 hours on a rotating device. Control samples were prepared with the same procedure without any digestive enzymes, adding MilliQ H_2_O instead. All the collected fractions were centrifuged to halt the enzymatic process (4 °C, 20 000 rcf·g, 10 min). Finally, 25 mL of supernatants were collected for further analysis, while pellets were freeze-dried and pooled to be used as pre-digested samples for the *in vitro* colonic fermentation.

### 2.4. Fecal Donors

Five healthy fecal donors were selected. The volunteers were 25-45 years of age, with a BMI of 18.5-25, non-smokers, and located in Wageningen (NL) and its surroundings. As exclusion criteria, all the volunteers must not have received antibiotic treatment 3 months before stool collection, must not have consumed pre or probiotic supplements before the experiment, and must not have a history of bowel disorders. Participants were informed about the study research, and their written consents were obtained. According to the Medical Ethical Advisory Committee of Wageningen University (METC-WU) guidelines, this research did not need ethical approval.

### 2.5. In-vitro batch fermentation

Colonic fermentation was carried out based on previously reported protocols, with some modifications ^23^. All the information relevant to in vitro fermentation has been included in the Supporting Information. With utmost brevity, a fecal microbiota supernatant (FMS) was prepared by homogenizing 40.0 g of fresh feces (five donors) in 200 mL of anaerobic phosphate buffer using a stomacher bag, and a buffered colon medium was prepared by mixing different amounts of K_2_HPO_4,_ NaHCO_3_, yeast extract, peptone, mucin, L-cysteine HCl, and Tween 80. Pre-digested samples from in vitro gastrointestinal digestion were mixed with buffered colon medium and FMS in 10 mL glass water-jacketed vessels. The vessels were placed in an incubator (37°C) on a rotating shaker (150 rpm). Batch cultures ran for a period of 48 hours, slurry fractions were taken at different time points (0, 6, 12, 24, and 48 hours); for donors 3, 4, and 5, the time point of 3 and 32 hours were also sampled. All slurry fractions were centrifuged and quenched with liquid N_2_ immediately after sampled and stored at −20 °C until further use. A total of 8 digestions in parallel were performed per donor: the three pre-digested burgers with FMS, the three pre-digested burgers without FMS (blank samples), and two blanks of the fermentation process, with and without FMS. More details about sample composition and assigned labels can be found in **Table S2**

### 2.6. Untargeted metabolomics approach

#### 2.6.1 Sample preparation

The extraction protocol was designed to obtain the non-polar and polar fractions of the samples coming from the *in vitro* gastrointestinal digestion (INFOGEST) and the colonic fermentation (C.F.). The current dual extraction method was adapted from previous studies that worked with fecal samples ^24–28^ The digestion sample flasks were thawed and vortexed for one minute, and then 100 μL of the homogenized sample was transferred into separate Eppendorf tubes. Sample was homogenized with 1 mL of MilliQ H_2_O: MeOH (1:1 (v/v)), and vortexed (90 s), followed by sonication (10 min) at low temperature. Then, a liquid-liquid extraction (LLE) was performed for the separation of the polar and the non-polar analytes by adding 600 μL MTBE. Additionally, 120μL of a polar mix of I.S. was added to each sample. Three cycles of vortex (30 s) and incubation in a refrigerator (5 minutes) were performed to ensure the proper transfer of the metabolites. Afterwards, the samples were centrifuged (10 min, 4°C, 12,557 rcf·g), and then, 300 μL of the upper organic phase and 1 mL of the polar fraction were transferred to glass vials. Glass vials with polar fractions were transferred to a Christ rotational vacuum concentrator to evaporate the solvents to dryness. For the evaporation of the organic solvent in the non-polar fractions, the vials were placed in a fume hood overnight, in the dark, and at room temperature. All vials were capped and stored at -80 °C before the reconstitution, the non-polar fraction was kept for lipidomics purposes. The polar fractions’ vials were reconstituted with 150 μL of MilliQ H_2_O: MeOH (9:1 v/v) in 0.1% F.A. Afterwards, vials were vortexed (90s), sonicated (10 min) and centrifuged (10 min, 4°C, 12,557 rcf·g). Finally, the samples were transferred into insert LC-MS vials, collecting 15 µL for a quality control (Q.C.) pool. Blank extraction samples were included following the same procedure. The instrument performance was monitored by using I.S. and Q.C.s.

#### 2.6.2 UHPLC-HRMS

The polar fraction analysis (5 μL) was carried out with a Nexera XS UHPLC system (Shimadzu Corporation, Kyoto, Japan) coupled to a LCMS-9030 quadrupole time-of-flight mass spectrometer (Shimadzu Corporation, Kyoto, Japan). The UHPLC unit consisted of a SIL-40C XS Autosampler, a LC-40D XS solvent delivery pump, a DGU-405 degassing unit, a CTO-40S column oven and a CBM-40 lite system controller. The qTOF-MS system was equipped with a standard electrospray ionization (ESI) source unit and a calibrant delivery system (CDS). The analysis was carried out in a Hydrophilic Interaction Liquid Chromatography (HILIC) and a reversed-phase liquid chromatography (R.P.) to analyse polar and medium polar compounds. A SeQuant® ZIC®-HILIC 5 μm particle size analytical column 150 x 4.6 mm (Merck KGaA, Darmstadt, Germany) and a Acquity UPLC BEH C18 column 1.7 μm particle sized analytical column 2.1 × 100 mm connected to an Acquity UPLC BEH C18 VanGuard Pre-column, 130Å, 1.7 µm, 2.1 mm ×5 mm (Waters Chromatography B.V., 4879 AH Etten-Leur, the Netherlands), were used for the HILIC and the R.P. analysis respectively.. The HILIC elution was performed using ACN (mobile phase A) and H_2_O (mobile phase B) in 0.1 % HCOOH and 10 mM NH_4_HCOO. The gradient started with 90% of A until 3.0 min, 70 % of A at 5.0 min, 20 % of A at 11 min, 20 % of A at 17 min, 90 % of A at 20 min with a total run time of 20 min and 0.7 mL min^-1^. For R.P. separation, H_2_O (mobile phase A) and MeOH (mobile phase B) both with 0.1 % HCOOH were used. The gradient started from 10 % B at 0 min to 90 % B at 14 min, 90 % B at 16 min, and 10 % B at 16.01 min with a total run of 18 min and a flow rate of 0.3 mL min^-1^. A total of 3 analysis were performed for both INFOGEST and C.F. including: HILIC and R.P. in positive ionization and R.P. in negative ionization. The column oven was set at 40 °C in all the acquisitions. Samples were randomly injected into the UHPLC-QTOF MS system with the aim of reducing the bias due to potential instrumental drift. The sample list was arranged as follows: the sequence begins with 5 extraction blanks, followed by 10 Q.C. injections that stabilize the column. Thereafter, one Q.C. and one extraction blank were injected into every ten and twenty samples, respectively.

The voltage of the ion-spray ionization was 4.0 kV and -3.0 kV for ESI+ and ESI-, respectively. For both negative and positive ionization, the heat block, desolvation line, and electrospray ionization probe temperatures were set at 400 °C, 250 °C, and 300 °C. At the same time, the flow rates were 2 mL min^-1^ for the nebulizer gas and 10 mL min^-1^ for the heating and drying gas. For data-dependent acquisition (DDA) analysis, MS^1^ and MS^2^ were acquired over an m/z range of 50-1000 Da and 100-1000Da, respectively. The number of DDA events was set to 18 with an event time of 0.05 s and a total loop time of 1 s, and the charge states were set between 1 and 3. Automatic exclusion of ions was performed by excluding other isotopes, ions of other charge states, and background artifact ions using a homemade exclusion list. The signal threshold was set at 1000, while the collision-induced dissociation (CID) energy ramp was set from 18 to 52 eV, with a gas pressure of 230 kPa. The mass axis and tune parameters were calibrated weekly with a sodium iodide (NaI) solution (Standard Sample for LCMS-9030, Shimadzu Corporation, Kyoto, Japan) . All the acquired mass spectra were internally calibrated by injecting ESI-L Low Concentration Tuning Mix (Agilent Technologies Netherlands BV, Amsterdam, Netherlands) through the sub-interface at the end of the run (equilibrating conditions).

#### 2.6.3 Data processing

UHPLC-QTOF MS raw data files were imported into MS-DIAL (MS-DIAL software, version: (MS-DIAL software, version: 4.9.2.2, Japan) for baseline filter, peak detection, retention time alignment, and annotation. Automated feature detection was performed from 0.5 min to 16 and 17 min for R.P. and HILIC, respectively. In all the analyses, mass signal extraction ranged from 50 to 1000 Da, while MS1 and MS2 tolerances were set to 0.01 and 0.03 Da, respectively. The minimum detection thresholds were set at 1000 and 10 for MS1 and MS2, respectively. For alignment purposes, a Q.C. from the middle of the sequence was selected as a reference file. The retention time tolerances were set at 0.2 and 0.5 min for R.P. and HILIC, respectively. After deconvolution, deconvoluted and annotated peaks were transferred to Excel to generate a matrix of *m/z* ratios, retention times, peak intensities, and MS/MS spectra of the deconvoluted ions. Features present in extraction blank, without MS/MS assigned, *m/z* match or MS/MS match were removed. In addition, those that showed low repeatability in the blanks of the fermentation process without fecal microbiota were discarded. Low repeatability was determined by using the RSD (>50%) and the average of the blank samples was more than one-tenth of the average of the feature in the groups with sample (R.M., PBCB and P.P.), to avoid discarding those features close to the limit of detection (LOD) in the group of blank samples.

MetaboAnalyst 5.0 was used to perform the data filtering, and statistics. To remove features with low repeatability or close to the baseline, features with RSD > 25% in Q.C. and interquartile range (IQR) < 40 % were discarded, respectively. Once the data set was filtered, the statistical study was focused on the metabolic variation between the two commercially available protein sources: R.M. and PBCB. For INFOGEST samples, one-factor statistical analysis was used to highlight the metabolic differences between both groups. In the case of colonic fermentation samples, it was interesting to explore these changes across the different time points of the colonic fermentation. Therefore, the option *time series + one factor* was selected for this purpose. Multivariate Empirical Bayes Time-series Analysis (MEBA) was used to rank features according to Hoteling’s T2 value, which indicates the differential temporal profiles. GraphPad Prism 10 (GraphPad Software, La Jolla, CA) was used to represent the results graphically. The findings were expressed using the mean values ± standard error of the mean of transformed data (square root, (SEM, N = 5) and differences were evaluated using two-ways repeated analysis of variance (ANOVA), considering a value of p < 0.05 statistically significant.

#### 2.6.4 Compound annotation

All openly accessible MS/MS spectral MSP-formatted libraries (MSPs) were used to tentatively annotate features during data processing. By the time this study was conducted (March 2023), these databases included 16,995 and 15,245 unique compounds for positive and negative ionization, respectively. Additionally, data processed with MS-DIAL, including unknown features, were exported to GNPS ^19^ for Feature-Based Molecular Networking (FBMN) ^29^ to enhance the annotation with GNPS spectral libraries ^30^. Those features that only present without MS/MS fragmentation were discarded.

The identification confidence level system introduced by Talavera Andújar et al. ^28^ was used to classify the features annotated by MS-DIAL. The *Dot product* parameter enables the assessment of the quality of match between the different samples and spectra in each MS/MS dataset. *In silico* fragmentation platform *MetFrag* ^31^ was used to confirm the MS/MS ion fragments matching. Those features whose annotations could not be explained were also discarded.

### 2.7 pH measuring

The pH measurement of all samples was performed using a pH electrode (VWR® PHenomenal® PH 1000 L). Before performing the measurement, the pH meter was calibrated. The process for preparing the sample involved mixing it in a vortex for 30 seconds before dissolving 0.5 mL of it in 4.5 mL of deionized water. The pH of the dilution was then measured.

### 2.8. Ammonia determination

A colorimetric assay kit (Ammonia Assay Kit AA0100, Sigma-Aldrich, St. Louis, USA) was used to analyze the ammonia content in the samples. To make sure the ammonia content was within the detection range of the kit, a 3x dilution factor with MilliQ H2O was used. Additionally, materials were clarified by centrifuging them (4°C, 12,557 rcf·g, 10 min) using an Eppendorf 5430R centrifuge. Then, a microplate spectrometer (SpectraMax ABS Plus, Molecular Devices, San Jose, USA) was used to measure the supernatant absorbance at 340 nm.

### 2.9. Indole analysis

A colorimetric assay kit determined total indole content (Indole Assay Kit MAK326, Sigma-Aldrich, St. Louis, USA). Before measurement, samples (150 μL) were diluted with 450 μL MilliQ H2O and centrifuged (4°C, 12,557 rcf·g, 10 min) using a 5430R Eppendorf centrifuge. 100 μL of the supernatant was used further for the analysis. The absorbance was measured at 565 nm using a microplate spectrometer (SpectraMax ABS Plus, Molecular Devices, San Jose, USA).

### 2.10. Phenol content

A colorimetric assay kit (Phenolic Compounds Assay Kit MAK365, Sigma-Aldrich, St. Louis, USA) determined the total phenol content. Pre-experiment data showed no dilution was needed to bring total phenol content to appropriate measuring ranges. Absorbance was measured at 480 nm using a microplate spectrometer (SpectraMax ABS Plus, Molecular Devices, San Jose, USA).

### 2.11 SCFAs analysis

The fermentation supernatants were subsequently centrifuged (4°C, 12,557 rcf·g, 5 min) and 2 mL were filtered (15 mm ∅, 0.2 µm regenerated cellulose filter, Phenomenex, Torrance, USA). Then, an internal standard of 2-ethylbutyric acid in 0.3 M HCl and 0.9 M oxalic acid was also added before injection into a gas chromatography system equipped with a flame ionization detector (GC-FID, GC-2014, Shimadzu, Hertogenbosch, Netherlands). The carrier gas employed was nitrogen. The temperature of ramp of GC-FID started at 100 °C, then increased to 180 °C for 2 min. Then, it increased to 240 °C at a rate 50 °C min^−1^ and was maintained at this temperature for 2 min. Standard calibration curves for acetic, propionic, butyric, valeric, iso-butyric, and isovaleric acids were created in the 0–50 mM concentration range.

## 3 RESULTS AND DISCUSION

### 3.1. *In vitro* bioavailability differences between beef and plant-based patties in the small intenstine

The present study used a simulated gastrointestinal digestion followed by a colonic fermentation model. The primary goal of the simulated gastrointestinal digestion was to provide a substrate for the subsequent colonic fermentation. By following the INFOGEST protocol ^22^, we initially analyzed the supernatant because it provides valuable information about compounds that are rapidly absorbed by intestinal epithelial cells in the small intestine. A total of 20 samples (supernatants) were collected at the end of the simulated gastrointestinal digestion, comprehending 5 replicas per patty, and 5 blanks. By employing untargeted metabolomics, we compared fingerprints of annotated metabolites that changed in response to the different feeding conditions. Firstly, principal component analysis (PCA) offered an exploratory visualization to observe trend groupings. **Figure S1** shows the PCA score plot obtained for the data sets corresponding to the polar fractions of gastrointestinal digestion. Q.C.s, a virtually “average” sample, are clustered in a central position, indicating minimal instrumental drift during acquisition. As observed, there is a clear separation between animal and plant-origin digested meats. Plant-based commercial and homemade plant-based burgers were clustered together in the score plot for both analyses (HILIC and R.P.) in the positive ionization mode, while for the R.P. analysis in the negative ionization mode there is a little but visible difference.

Univariate and multivariate statistical analyses were then applied to investigate the metabolic differences between the different feeding conditions. Only those features that met the following characteristics were chosen t-test (p-value<0.05), volcano plot (p-value<0.05), fold-change>2, OPLS-DA (pcorr>|0.8| and p[1]>|0.1|). The selected markers, including 32 markers of RM and 45 markers of PBCB, are recorded in **Table S3.** Among the features highlighted for beef meat digested samples, it is noteworthy to mention the presence of carnitine and acylcarnitines such as acetylcarnitine, butyryl carnitine, propionyl carnitine, hexanoyl-L-carnitine. Carnitine and derivates are known biomarkers of beef consumption, and they have essential roles in energy production by lipid β-oxidation in the mitochondria of muscle cells in mammals ^32^. Other statistically known red meat biomarkers found in beef patties were L-carnosine, L-anserine, taurine, creatine, and acylcholine ^33^. Triterpene and steroidal saponins, triterpenoids, oligosaccharides, flavones, lignans, and purine and purine derivates were the compounds found in the intestinal fraction of the digested plant-based commercial analog, which are compounds naturally found in pea beans ^34^.

We also noticed that stachyose, an α-galactoside known in pea beans, presents a high resistance to hydrolysis by digestive enzymes and was at higher abundance after performing the *in vitro* digestion. *In vivo*, this α-galactoside is not absorbed in the upper part of the gastrointestinal tract. Stachyose is known to reach the large bowel and be used as a substrate by several gut bacteria ^35^ highlighting an *in vitro* modeling limitation and, at the same time, untargeted metabolomics demonstrated to be a reliable tool to screen bioavailable nutrients and leftovers reaching the colon. Among the amino acids found in R.M alanine, glutamine, leucine, lysine, methionine, phenylalanine, tryptophan, and valine were the most abundant, while proline and asparagine were more predominant in PBCB. In addition, differences in the levels of some peptides (generally di and tri-peptides) were found. *In vivo* studies on mice also pointed out variations in the peptide composition resulting from the digestion of both types of burgers ^36^. According to the food matrix characterization performed by ^37^, the average amino acid composition in plant-based burgers (pea and soy) is quite similar to meat-based burgers. Plant-based alternatives presented low methionine, glycine, and lysine levels but high glutamic and aspartic acid levels. The findings of the current study suggest that the composition of the food matrix potentially affects amino acid digestibility, which is generally higher in burgers of animal origin. In this regard, Xie et al. showed that consuming animal-based burgers increased the transfer of essential and non-essential amino acids from the jejunum to the bloodstream compared to the digestion of their PBMA counterpart ^36^.

### 3.2. Modeling the impact of undigested proteins in the colonic region

The undigested fraction (pellet) generated during the last stage of the simulated gastrointestinal digestion was employed as substrate for the *in vitro* colonic fermentation model, collecting 233 samples longitudinally. **Table S4** compiles the final list of samples collected.

#### 3.2.1. A rapid biochemical profiling comparing red meat and meat analogs

##### 3.2.1.1 Ammonia

Ammonia, consisting of NH_4_ ^+^ and NH is produced by bacterial deamination of amino acids ^38^. It is a proteolysis marker that is highly sensitive to changes in the fermentation of amino acids. The detrimental effects of high elevated levels of ammonia on the large intestine have been extensively studied in the past, including the production of histological damage to the colonic mucus layer, the reduction of absorptive capacity and lifespan of colonocytes, the increase in the expression of marker genes reflecting mucosal cell turnover and the synthesis of pro-inflammatory cytokines ^9^. In the current study, ammonia levels increase for all protein sources during the first 6 hours to remain stable the remainder of the fermentation (**Fig. 1A**). The R.M. had the highest ammonia concentration at all time-points, except at the 6 h (**Fig. 1A**). Smith & Macfarlane (1997) compared the dissimilation of 20 amino acids by human fecal microbiota and found that arginine, lysine, histidine, and glycine resulted in the highest ammonia production in a 24-hour fermentation period ^39^. According to De Marchi et al., 2021, meat-based burgers contain more glycine than plant-based burgers, which might explain the consistently higher ammonia concentration in the samples ^37^. In addition, compared with real meat, plant-based meat analogues were less digested and released fewer peptides, which downregulated the gene expression involved in gastrointestinal nitrogen nutrient sensors, and further reduced the gastrointestinal digestive function ^36^. In this case, more protein might enter large intestine in PBCB group compared with R.M. group. However, there were little differences in ammonia content between R.M. and PBCB in this research work. One reason for the nonsignificant correlation might be the short-term intervention due to the *in vitro* model. Thus, it is worthwhile to focus on this point in the case of long-term *in vivo* experiments.

**Figure 1.**
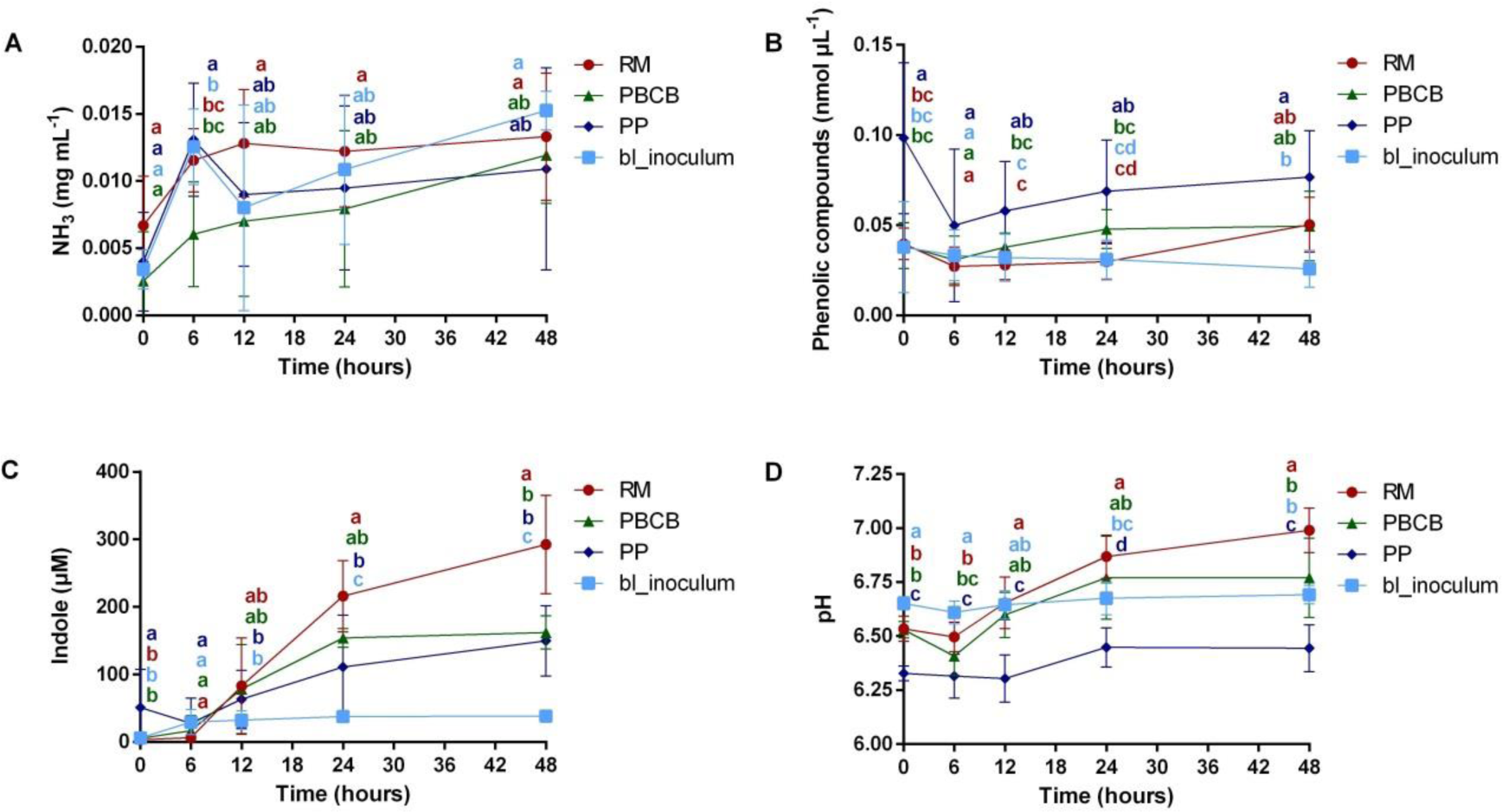
The changes in ammonia (a), total phenolic compounds (b), total indoles (c), and pH (d) during the in vitro colonic fermentations. Blank inoculum, R.M., PBCB, and P.P. samples are depicted. The different letters indicated the significant difference by One-way ANOVA analysis for different groups at the same time point.

##### 3.2.1.2 Phenols

Degradation products from tyrosine include 4-hydroxyphenylpyruvate, 4-hydroxyphenyllactate, 4-hydroxyphenylpropionate and 4-hydroxyphenylacetate as well as phenol, p-cresol and 4-ethylphenol. Phenylalanine bacterial metabolism leads to similar derivatives ^40^. The total phenol content of P.P. was higher than all other samples at all time points, with a statistically significant difference (p<0.05) at time-point 0 (**Fig. 1B**). It is interesting to note that R.M., PBCB, and the blank inoculum show the same levels at time 0. In contrast, P.P. offers a considerably higher concentration of phenols, most likely due to the presence of polyphenols. This difference may also be attributed to the amino acid composition and digestibility of proteins between P.P. and R.M. During fermentation, phenol content in R.M. group increased, which is mainly due to the production of tyrosine by microbial catabolism. This is supported by the amino acid content result showing that tyrosine gradually decreased during fermentation period (**Fig. 3C**). Finally, there is little difference in phenolic compounds content between R.M. and PBCB groups after 48 h of fermentation. Polyphenols have been linked with beneficial action on multiple disorders, including obesity, diabetes and cardiometabolic and neurodegenerative diseases, which might be due to their antioxidant and anti-inflammatory effects. Evidence suggests that polyphenols are able to express prebiotic properties and exert antimicrobial activities against pathogenic gut microflora ^41^. However, viability of colonic epithelial cells isolated from human biopsies was decreased after exposure to 1.25 mM phenol, a physiologically relevant concentration, whereas clearly higher phenol concentrations (20 mM) were required to reduce viability of HT-29 cell ^40^. Therefore, phenolic compounds perform either beneficial or deleterious effects on colon, and this is largely dependent on their luminal concentrations and the colonic microenvironment of host individuals. In vivo, these phenolic compounds would be largely absorbed from the colon, detoxified in the colon mucosa and the liver by glucuronide and sulfate conjugation, and finally excreted in urine ^40^.

##### 3.2.1.3 Indoles

Indoles are produced by microbial conversion of tryptophan via the enzyme tryptophanase, exclusively found in bacteria ^42^. In this section, we quantified the sum of indoles, and indole concentrations increased over time (**Fig 1C**). Within the first 12 h of fermentation, protein samples had similar indole content, with almost no difference observed between the R.M. and PBCB samples. However, after 12 h of fermentation, the indole production increased considerably, especially in the R.M. sample. The R.M. group had a significantly higher indole concentration by the end of the fermentation time. On the one hand, indoles increase the expression of tight-junction proteins in the gut epithelium, improve intestinal barrier function, and attenuate inflammatory markers ^43^. On the other hand, high concentrations of indoles thought enzymatic transformations in the liver and colonocytes might result in indoxyl sulfate, which is considered a uremic toxin ^44^. In this *in vitro* work, the R.M. group exhibited a significantly higher fecal indole content than PBCB and P.P. groups, suggesting that beef proteins may increase indole production. This is consistent with the amino acid content result showing that tryptophan largely decreased in R.M. group during fermentation period (**Fig. 3D**).

##### 3.2.1.4 pH

As a general trend, the pH increased with the fermentation time (**Fig. 1D**). In the colon, the main contributors that modulate pH are molecular hydrogen, bicarbonate, organic acids, ammonia, and other microbial metabolites ^45^. It is tempting to suggest that the increase in pH for the samples is correlated with the increase in ammonia. Generally, a lower colonic pH is associated with better colon health since it could inhibit the proliferation of undesirable pathogens and affect microbial enzymes’ activity ^46^. In this respect, the findings of this study suggest that plant-based proteins could generate a lower pH than meat proteins.

##### 3.2.1.5 SCFA

Although SCFA are the major end products from carbohydrate fermentation, they are also produced from many amino acids by reductive deamination ^40^. BCFA (e.g., isobutyric acid and isovaleric acid) are produced in the gut upon proteolytic fermentation of branched-chain amino acids (leucine, valine, and isoleucine) ^47^. SCFA are rapidly absorbed from the colon and are generally considered to be beneficial for the host. Among other biological roles SCFAs provide energy for intestinal epithelial cells and peripheral tissue, inhibiting potential pathogen growth, stimulating epithelial proliferation, facilitating tight junction formation, and inhibiting inflammation and genotoxicity ^9^. In this work, the acetic acid content increased with fermentation time for R.M., PBCB, and P.P. samples (**Fig. 2A**), showing a significant difference in R.M. group at time point 24h. In the case of propionic acid, no significant differences were observed between the three groups. The iso butyric acid content increased after 12 h for R.M., PBCB, and P.P. samples, and the significant difference among the three groups occurs from the time point 24 h, at which the iso butyric acid content in R.M. is higher than that of PBCB and P.P. This larger amount of BCFA in R. M. group might be related to the higher leucine and valine contents in R.M. matrix (as mentioned above). In addition, butyric acid content increased for R.M. and PBCB groups, but there was no noticeable change in the P.P. group. Butyrate is the most important energy source for colonocytes and plays a major role in proliferation and differentiation ^48^. In vivo, butyric acid may have improved gut barrier integrity (Fan et al., 2023). The fecal butyric acid might be increased by increasing the abundance of *Lactobacillus reuteri, Lactobacillus spp., Lactobacillus lactis* and *Lactobacillus plantarum* ^50^. A similar trend was also seen for isovaleric acid. According to these results, different protein sources distinctly altered the SCFA production.

**Figure 2.**
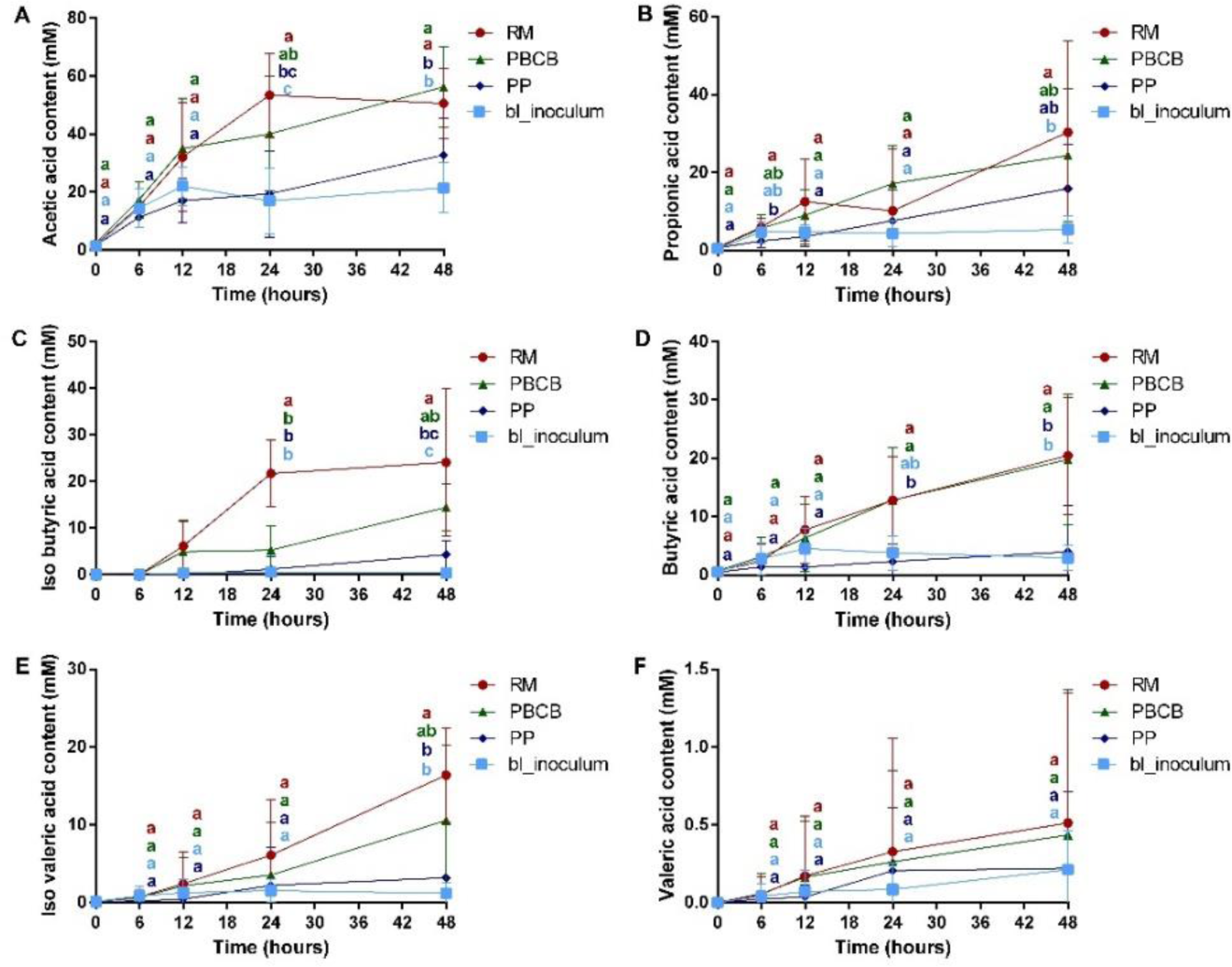
The changes in SCFAs contents during fermentation for blank inoculum, R.M., PBCB, and P.P. samples. (A-F) the significant difference analysis for different samples at the same time point. The different letters indicated the significant difference by two-way ANOVA analysis for different groups at the same time point.

**Figure 3.**
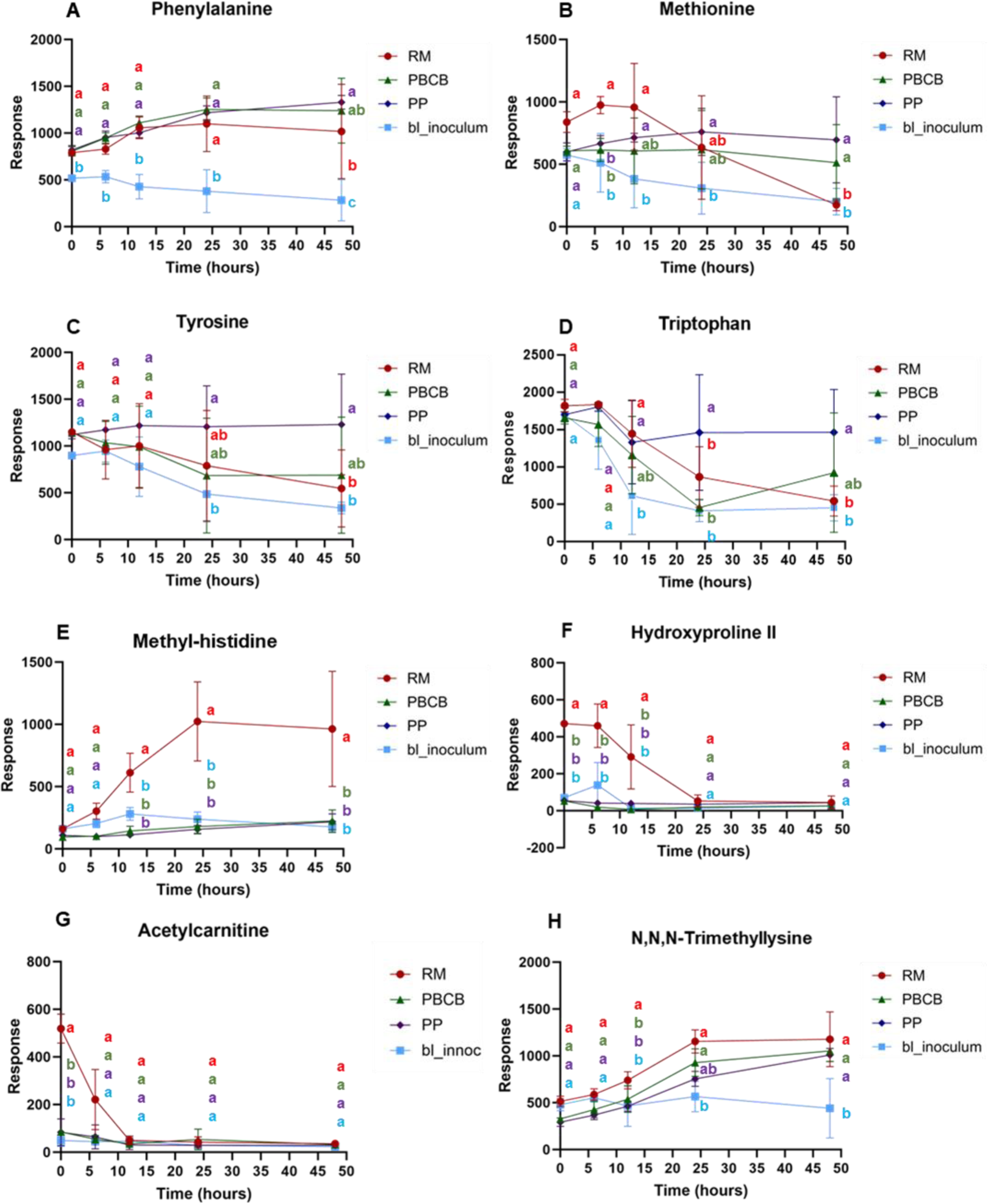
Response of (A) phenylalanine, (B) methionine, (C) tyrosine, (D) tryptophan, (E) methyl-histidine, (F) hydroxyproline II, and (G) acetylcarnitine, (H) N,N,N-Trimethyllysine. Notes: RM beef burger, PBCB plant-based commercial burger, PP plant-based homemade burger, bl_inoculum, sample without any substrate. The different letters indicated the significant difference by two-way ANOVA analysis for different groups at the same time point.

#### 3.2.2 Protein-related GMMs elucidated by an untargeted metabolomics workflow

Samples were extracted and analyzed following the untargeted metabolomics approach designed for polar and semi-polar metabolites (*see section 2.6.1*). After data processing, annotation, and data filtering, a total of 343 features for HILIC ESI (+), 306 for RP ESI (+), and 157 for RP ESI (-) were collected. Q.C. pool grouping in the PCA score plot (**Fig. S2**) indicates adequate analytical performance across all three acquisitions. As shown in **Figure S2**, the highest degree of dispersion was given within those groups containing undigested substrate and fecal inoculum (R.M., PBCB, and P.P.). However, PCA or supervised multivariate models as partial least squares discriminant analysis (PLS-DA) are inadequate models to evaluate the metabolic changes in a time-series metabolomics, in contrast to alternative approaches like MEBA, which provides a valuable tool for assessing the variability of both, within and between time points ^51,52^. This approach was used to rank the potential metabolites of interest according to their Hoteling’s T2 value by comparing R.M. and PBCB groups across the colonic fermentation and selecting only those features with a Hoteling T2 > 20. Special attention was paid to the removing of non-informative markers before modeling the data. First, those features showed low repeatability in both Q.C. replicates and blanks of the fermentation process without fecal inoculum. Second, variables that were almost constant across the experiment conditions and those with a signal close to the baseline. **Table S5** collects all the selected annotated features through the untargeted metabolomics workflow sorted out by ontology together with the level of identification. Among the families of compounds highlighted through the metabolomics pipeline were amino acids and derivates, acetylated amino acids, dipeptides, and hybrid peptides. The combination of R.P. LC and HILIC was useful to cover a broader range of polarities and expand the metabolic scope of the study. For example, the elucidation of highly polar compounds, such as acetylated amino acids, would not have been possible without using HILIC. The following sections describe the trends observed for every chemical family and discuss their potential implications on gut health.

##### 3.2.2.1 Amino acids and derivates

Among all the amino acids annotated, only phenylalanine, methionine, tyrosine, and tryptophan (**Fig 3, A, B, C, and D**, respectively) showed notable differences between RM and PBCB during the colonic fermentation. It is worth noting that the inoculated samples without substrate (bl_inoculum) presented high basal levels of amino acids due to the medium composition (*see section 2.5*). Phenylalanine **(Fig. 3, A)** was released during colonic fermentation in beef and pea-based meats, with slightly higher levels in the second group. In the case of methionine **(Fig 3, B)**, as expected, RM stood out from the rest at basal levels, experiencing, at 12 hours, more accelerated catabolism than the rest of the groups. Higher levels of tyrosine **(Fig 3, C)** and tryptophan **(Fig 3, D)** were observed in the PP than in the rest of the groups throughout the colonic fermentation. Phenylalanine, tryptophan, and histidine metabolomic pathways are closely related to gut inflammation ^53^. In the case of phenylalanine and tryptophan, no relevant derived metabolites were found through the metabolomics pipeline. However, the histidine derivative methyl-histidine **(Fig 3, E)** was discovered and showed a different trend in beef burger samples released during the first 24 hours of the fermentation. Even though histidine was not highlighted in the MEBA analysis (**Fig S3, A**), its methylation can occur at either the N1 or N3 position of its imidazole ring, resulting in 1-methyl (1-MH) and 3-methyl histidine (3-MH) ^54^. Both isomers, 1-MH and 3-MH, can only be produced from histidine residues by methyltransferases in mammals ^55^. 3-MH, as a part of the histidine metabolic pathway, may have a role in inflammation diseases. However, the association was not established in stool on a mice model^53^. An *in vivo* study analyzing feces has recently found lower amounts of 3-MH and anserine (*see section 3.2.2*) in patients with irritable bowel syndrome ^56^.

Regarding the proline derivatives, there were two features annotated as hydroxyproline: hydroxyproline I, measured in RP ESI (+), and hydroxyproline II in HILIC ESI (+). In both cases, two crucial structural fragment ions were found: *m/z* 86.0606 and *m/z* 68.0500, both associated with the pyrrolidine structure of the proline but with different trends observed. Hydroxyproline I (**Fig S3, b**) showed the same basal level for all the groups. At the same time, a significant amount of hydroxyproline II (Fig 3, F) was observed in RM. Hydroxyproline I increased during the first 12 hours of the fermentation. By contrast, hydroxyproline II may have been catabolized by the gut microbiota. Hydroxyproline II may correspond to 4-hydroxyproline, a known biomarker of red meat that is a major component of protein collagen and plays a key role in collagen stability. This proline-derivative is known for improving bone, joint, and skin stability, and some animal models reveal its role in preventing gut inflammation ^57^. Studies in rats claimed that most of the 4-hydroxyproline found in blood comes from free 4-hydroxyproline and 4-hydroxyproline-containing di- and tri-peptides whose hydrolysis takes place in the enterocytes ^58^, but there is no evidence about being anabolized by gut microbiota. Although 4-hydroxyproline is known to play important roles in the metabolic pathway for glycine production, little is known about its catabolism. The findings of our study reveal the potential role of gut microbiota in releasing 4-hydroxyproline and open the door to further research in this regard.

L-acetylcarnitine was found at high levels in RM samples at time 0h, and it is consumed abruptly during colonic fermentation (**Fig 3, G**). In chemical terms, acetylcarnitine corresponds to the acetylated derivate of the amino acid L-carnitine and is acquired by ingesting meat of animal origin (*see Section 3.1*). L-acetylcarnitine role is often associated with controlling energy metabolism in the mitochondria, but it also plays a role as a substrate of the carnitine pathway. The final product of this metabolic pathway is trimethylamine (TMN), which is known to be transformed into trimethylamine N-oxide (TMAO) in the liver. High TMAO levels have been associated with an increased risk of cardiovascular disease ^32,59^. Unfortunately, due to the low molecular mass of TMN, it has not been possible to identify it through our untargeted metabolomic approach. Note that despite the absence of acetylcarnitine/carnitine in plant-based meat substitutes, trimethyllysine (**Fig 3, H**), a component of the carnitine metabolic pathway, can be generated during fermentation.

For all the groups under study, N,N,N-trimethyllysine begins at the same basal level and is increased during the fermentation of both animal and plant-based patties. The increase is more pronounced in the RM samples in the first 12 hours of fermentation, although these levels are comparable to those of plant-based patties after 24 hours. N,N,N-trimethyllysine is formed by protein lysine methylation, particularly histone proteins, and it plays an important role in carnitine biosynthesis and epigenetics ^60^. However, *in vivo* studies established associations between high levels of N,N,N-trimethyllysine in serum and the presence and severity of heart failure related to the carnitine pathway ^61^. To our knowledge, this is the first time there is evidence of the anabolism of trimethyllysine during colonic fermentation. The hydrolysis of dietary methylated proteins may produce trimethyllysine ^61^.

**Fig. 4** shows the levels of four N-acyl amides annotated by the metabolomics workflow. N-acyl amides are a form of endogenous fatty acid molecule composed of a fatty acyl group linked to a primary amine metabolite via an amide bond. The N-acyl amides identified are phenylalanine linkages with caprylic, linoleic, and α-linoleic acid **(Fig 4, A, B, and C, respectively**), and leucine with linoleic acid **(Fig 4, D)**. In all four cases, a precise release of these compounds is observed in the PBCB group during colonic fermentation. This phenomenon is not so evident in the PP group, except for N-caprylyl-L-phenylalanine **(Fig 4, A)**, where the production of this N-acyl amide is even higher. The availability of fatty acids could condition the production of N-acyl amides. Although PP patties were designed to mimic PBCB, there are differences in the source of fat. PBCB ingredients included coconut and canola oil, while PP only had coconut oil. Canola oil is richer in linoleic and α-linoleic acid but poorer in caprylic acid ^62^. Also, caprylic, linoleic, and α-linoleic acid levels are higher in plant-based commercial burgers than meat-based burgers ^37^. The gut microbiota produces n-acyl amides, and it has been noted that these substances are crucial for controlling the physiology of the gastrointestinal tract ^63^. As a result, increased levels of these compounds may positively affect gut health.

**Figure 4.**
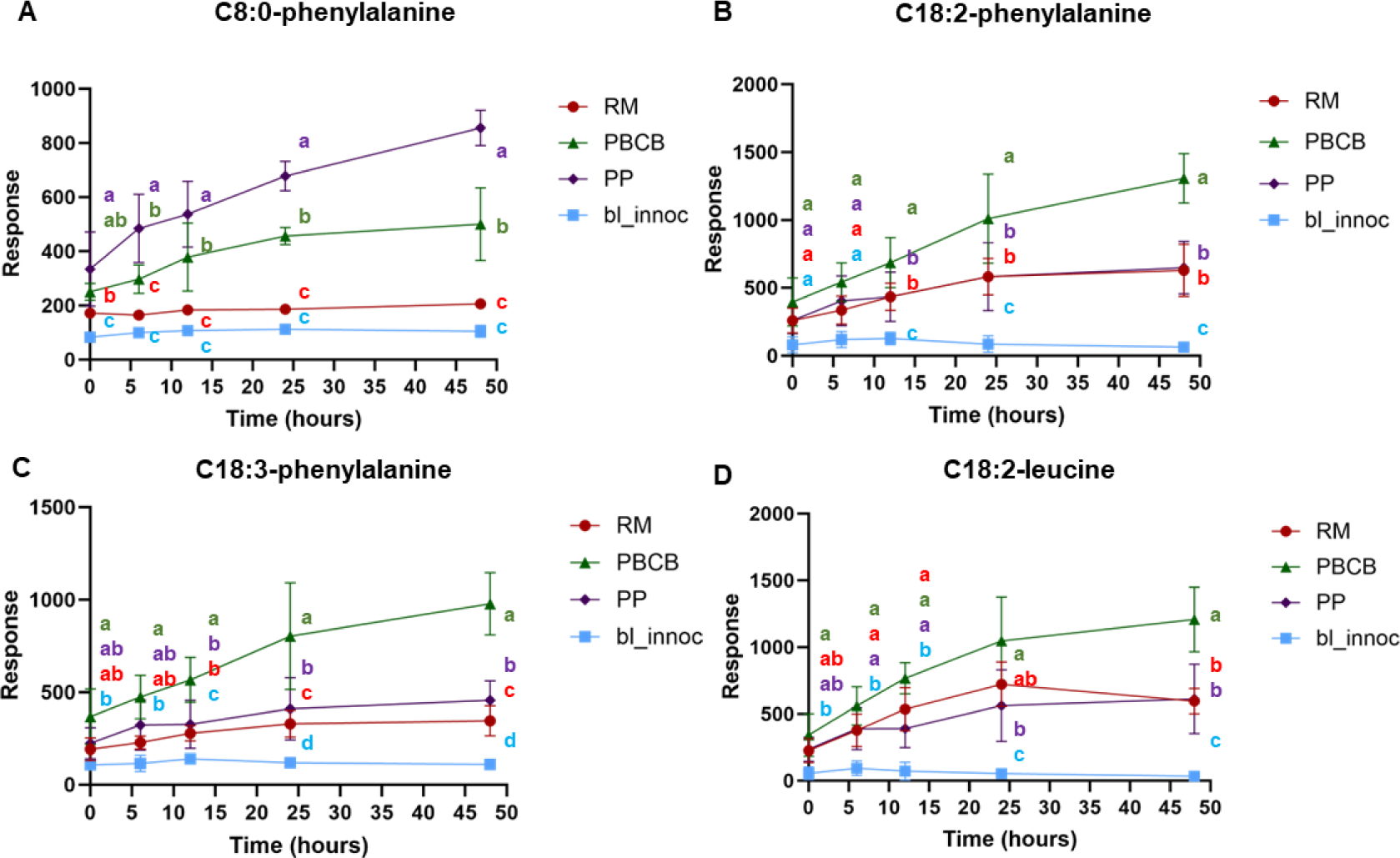
Response of (A) C8:0-phenylalanine, (B) C18:2-phenylalanine, (C) C18:3-pheynlalanine, (D) C18:2-leucine. Notes Notes: R.M. beef burger, PBCB plant-based commercial burger, P.P. plant-based homemade burger, bl_inoculum, sample without any substrate. The different letters indicated the significant difference by two-way ANOVA analysis for different groups at the same time point.

#### 3.2.2.1 Dipeptides and derivative

Several dipeptides have been highlighted in the elucidation workflow **(Fig 5.)**. Two main trends are observed. (i) Most of them presented high levels induced by the substrate at the beginning of the fermentation, except for the high basal levels of pyroglutamic-(iso)leucine, which is microbial-derived. (ii) The gut microbiota catabolized all of them during the first 12-24 hours of fermentation. Arginine-(iso)leucine, lysine-(iso)leucine, and pyroglutamyl-(iso)leucine (**Fig 5. B, E, and F**), presented the same levels at time 0 across all feeding conditions, while alanine-(iso)leucine, histidine-(iso)leucine, (**Fig 5. A and C**) showed a higher level on RM group. On the contrary, phenylalanine-arginine (**Fig 5. G**) was only found in plant-based alternatives, while isoleucine-methionine (**Fig 5. D**) is only present in beef meat. The same trend was observed for the hybrid peptides L-carnosine (**Fig 6. A**) and, at a slower rate, not for all donors, for L-anserine (**Fig 6. B**). L-carnosine has been found to potentially have an antioxidative role in intestinal epithelial cells ^64^, and its supplementation has been associated with the reduction of glucose, blood pressure, and obesity in rats with metabolic syndrome ^65^. Previous studies have reported decreased di- and tripeptides along the colon during digestion ^66^. In the case of anserine, a dipeptide containing beta-alanine and 3-methyl-histidine, a study in mice reveals its role in an important physiological function in intestinal health, including prevention of gut microbiota dysbiosis, repair of the intestinal epithelial barrier and production of SCFAs ^67^. Peptides derived from protein digestion are thought to have significant functions in transmitting signals to gastrointestinal cells, stimulating intestinal tract ^68^. According to a recent study on the effects of PBMA intake on mice’s gastrointestinal digestive function, ingesting PBMA causes less peptide production along the small intestine, and gastrointestinal digestion is less efficient than eating actual meat ^36^. The findings of the current research, considering the limitation of *in vitro* digestion models, showed comparable trends in both matrices regarding the transfer of dipeptides from the small intestine to the colonic stage and in their subsequent catabolism by the action of the microbiota.

**Figure 5.**
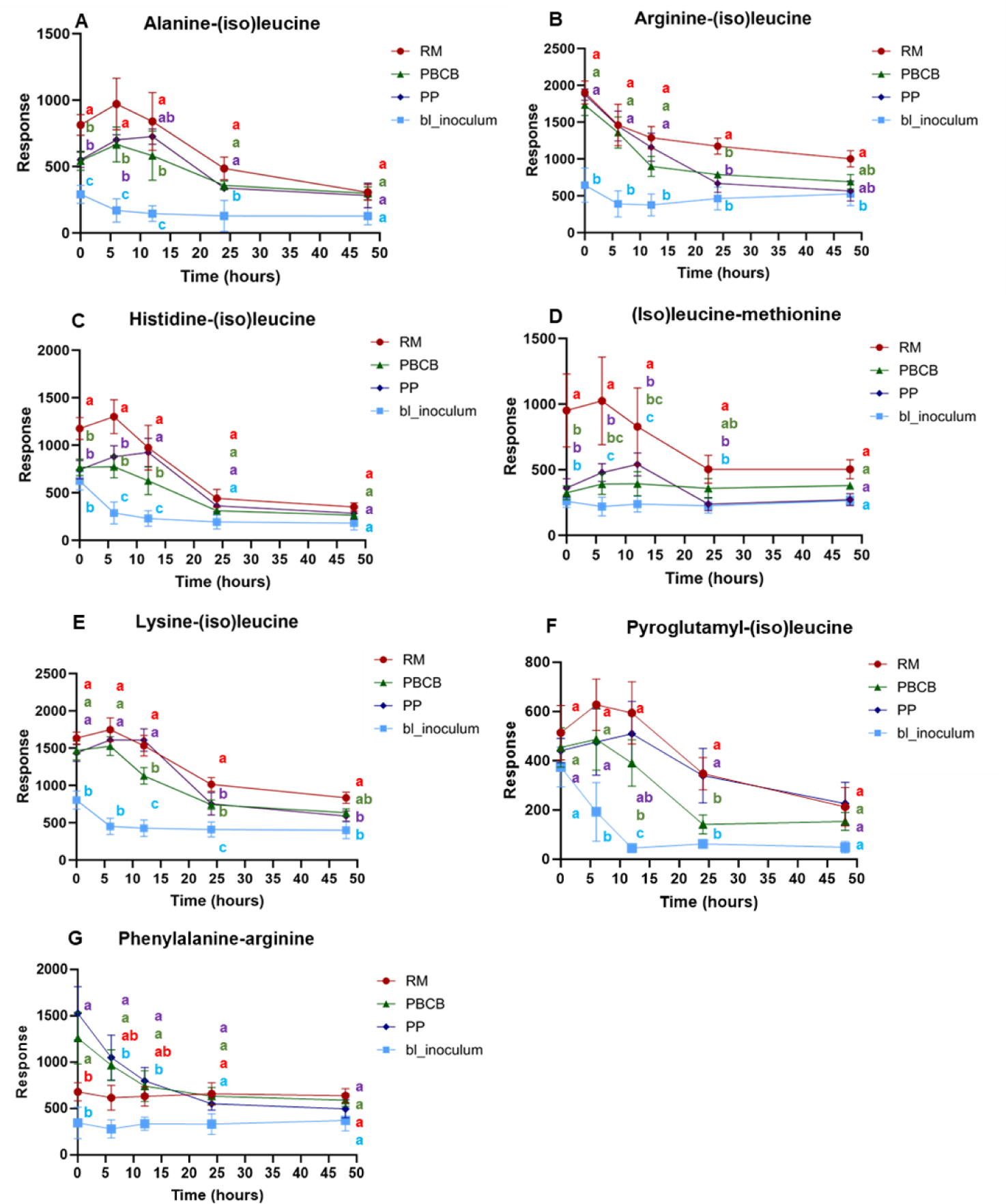
Response of (A) alanine-(iso)leucine, (B) arginine-(iso)leucine, (C) histidine-(iso)leucine, (D) (iso)leucine-methionine, (E) lysine-(iso)leucine), (F) pyroglutamyl-(iso)leucine and (G) phenylalanine-arginine. Notes: R.M. beef burger, PBCB plant-based commercial burger, P.P. plant-based homemade burger, bl_inoculum, sample without any substrate. The different letters indicated the significant difference by two-way ANOVA analysis for different groups at the same time point.

**Figure 6.**
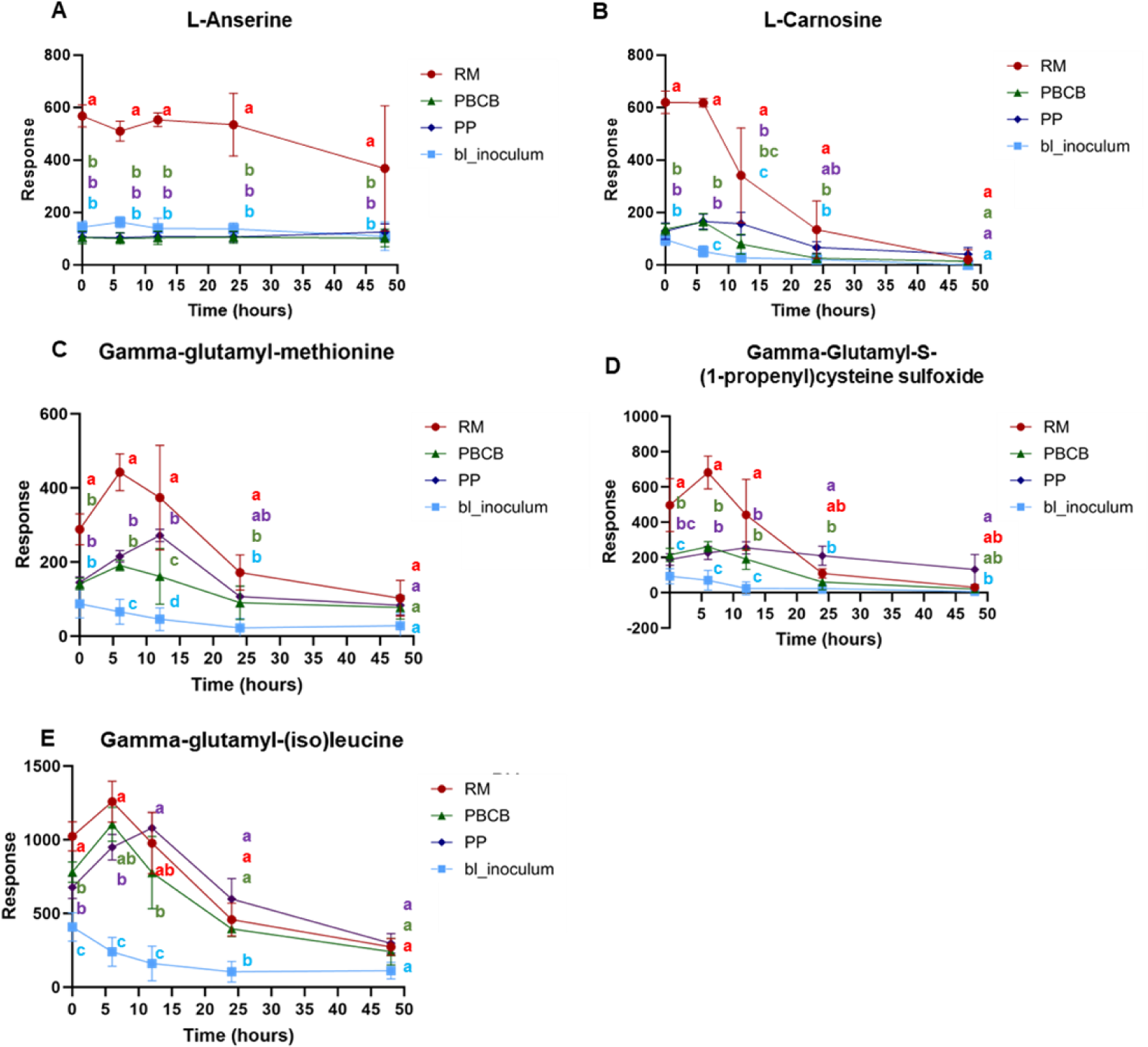
Response of (A) L-anserine, (B) L-carnosine, (C) gamma-glutamyl-methionine, (D) gamma-glutamyl-S-(1-propenyl)cysteine sulfoxide, (E) gamma-glutamyl-(iso)leucine. Notes: R.M. beef burger, PBCB plant-based commercial burger, P.P. plant-based homemade burger, bl_inoculum, sample without any substrate. The different letters indicated the significant difference by two-way ANOVA analysis for different groups at the same time point.

Our study observed that gut microbiota was also capable of releasing some dipeptides, particularly gamma-glutamyl amino acids. Gamma-glutamyl leucine (**Fig 6. C**) was released during the first stages of the colonic fermentation and then rapidly utilized by the microbial communities. This trend was observed in the three treatments, but it was more noticeable in beef meat since the basal levels of this compound were also higher in the animal-based protein. A similar up-down trend was observed for gamma-glutamyl methionine and gamma-glutamyl-S-(1-propenyl) cysteine sulfoxide (**Fig 6. D y E**) on R.M. but not in plant analogs. Gamma-glutamyl transpeptidase (GGT) is a group of enzymes that catalyzes the transfer of the gamma-glutamyl group from gamma-glutamyl peptides to other peptides or free amino acids, and it is believed to be responsible for many physiological disorders in mammals, such as oxidative stress and gut inflammation ^56^. In humans, these enzymes are primarily present in biliary epithelial cells but can also be present in prokaryotic cells ^69^. In all the groups studied, gut microbiota can reverse gamma-glutamyl amino acids to baseline; however, epithelial cells could absorb these compounds in the early stages of colonic fermentation. In this hypothetical case, further research is necessary to investigate the potential effects of gamma-glutamyl-methionine intake and gamma-glutamyl-S-(1-propenyl) cysteine in human health.

## 4 CONCLUSIONS

Our *in-vitro* approaches provided evidence that the digestion of a pea-based PBMA compared to a beef burger causes alterations in protein digestibility and the bioavailability of their associated metabolites. Evaluation of the rapidly absorbed metabolite-rich fraction in gastrointestinal digestion using an untargeted metabolomics approach revealed differences in the composition of small peptides and increased bioavailability of free amino acids in animal-based burgers. During the *in vitro* colonic fermentation, no differences in ammonia content were observed between R.M. and PBCB. In the case of the total phenol content, P.P. showed higher basal concentrations than the rest of the groups, although phenol content in R.M. group largely increased due to the production of tyrosine by microbial catabolism. R.M. samples exhibited a significantly higher indole content than the plant-derived samples. In terms of pH, as a general trend it increased during the fermentation for all groups, although lower levels were observed for plant-derived patties. For SCFAs, after 24 h of fermentation, the acetic acid content in R.M. is higher than that of other groups. The significant difference in iso butyric acid content among the three groups occurs from the time point 24 h, at which the iso butyric acid content in R.M. is higher than that of PBCB and P.P. In addition, butyric acid content increased for R.M. and PBCB groups during fermentation. The untargeted metabolomics approach on *in vitro* colonic fermentation samples revealed different trends in protein-related amino acids between beef-based and pea-based proteins, with potential implications for gut health. Using real meat as a feeding condition, we noticed anabolism of metabolites related to gut inflammation, such as 3-methyl-histidine, part of the histidine pathway, and gamma-glutamyl amino acids. However, the intake of beef proteins also incorporates metabolites with essential physiological functions for the gut microbiome, such as 4-hydroxyproline, carnosine, and anserine, which are catabolized by the microbiota during colonic fermentation and not present in pea proteins. In addition, the list of meat-unique compounds also includes acetylcarnitine, whose role in gut health is controversial as the carnitine pathway has been linked to cardiovascular disease. Related to carnitine metabolism, the findings of our study revealed that gut microbiota can release trimethyl lysine. Although it is more acute in meat patties, plant-based alternatives could also release this precursor of the carnitine pathway. PBMA patties, overall PBCB, showed a high production of N-acyl amino acids during colonic fermentation compared with real meat, which are associated with signaling functions in the intestine tract. Finally, and related to signaling functions in the gut, in both animal and plant-based burgers, high levels of dipeptides have been found at the beginning of colonic fermentation, which were catabolized after 24 hours. Generally, the levels of dipeptides were more elevated in real meat, although there are some exceptions, such as phenylalanine-arginine.

## ACKNOWLEDGEMENTS

1. D. Izquierdo-Sandoval acknowledges the Ministry of Science, Innovation and Universities of Spain for funding his research through the FPU pre-doctoral program (FPU19/ 01839) and for the financial support received for his research stay at the Wageningen University & Research. T. Portoles acknowledges Ramon y Cajal Program from the Ministry of Economy and Competitiveness, Spain (RYC-2017-22525) for funding her research.

