## Supplemental Material for "In-vitro digestion models to examine the effects of Plant-Based Meat substitutes on Gut Microbial Metabolites"

### Supplementary Material

**An *in vitro* modeling approach comparing plant-based analogs and red meat reveals differences in gut microbial metabolite profiling.**

David Izquierdo-Sandoval[1], Xiang Duan[2,3], Christos Fryganas[3], Tania Portolés[1], Juan Vicente Sancho[1], Josep Rubert[3,4*]

[1] Enviromental and Public Health Analytical Chemistry, Research Institute for Pesticides and Water (IUPA), Universitat Jaume I, Av. Sos Baynat S/N, 12071 Castellón de la Plana, Spain.

[2] College of Food Science and Engineering, Northwest A&F University, Yangling 712100, PR China

[3] Food Quality and Design, Wageningen University & Research, Bornse Weilanden 9, Wageningen, 6708 WG, Netherlands.

[4] Division of Human Nutrition and Health, Wageningen University & Research, Stippeneng 4, Wageningen, 6708 WE, Netherlands.

**Corresponding author:**

Josep Rubert, PhD

Assistant Professor in Gastrointestinal Health

Wageningen University & Research

Division of Human Nutrition and Health (Nutritional Biology) & Food Quality and Design.

P.O. Box 17, 6700 AA Wageningen. Wageningen Campus,

HELIX - Building Stippeneng 4 | 6708 WE Wageningen

T: +31 (0) 317 486 842 | M: +31 (0) 618 520 624 | E:

**TABLE OF CONTENTS:**

Title page........................................................................................................................S-1 - S-3

Material and methods.......................................................................................................S-5

Simulated in vitro gastrointestinal digestion..........................................................................S-5

In-vitro batch fermentation.................................................................................................S-5

Table S1. Composition of foodstuff material....................................................................................S-6

Table S2. Composition of the flasks subjected to the in vitro colonic fermentation. Sample labelling is in parentheses next to the treatment name......................................................................................S-6

Table S3. Features of interest obtained in simulated "in vitro" gastrointestinal digestion. The feature is defined as "retention time (min)" _ "m/z". It includes the name of the compound, the formula, group, INCHIKEY, the ontology and the level of identification…......................................S-7 - S-14

Table S4. Samples collected during the colonic fermentation. It includes sample name, class, time, and subject...........................................................................................................S-15 – S-21

Table S5. Features of interest obtained in colonic fermentation. The feature is defined as "retention time (min)" _ "m/z". It includes the name of the compound, the formula, the INCHIKEY, the ontology and the level of identification..............................................................................S-22 – S-23

Figure S1. PCA score plot component 1 vs component 2 for the simulated gastrointestinal digestion metabolic profiles, mirroring the small intestine content. Four groups of samples were analyzed from this in vitro digestion model: Beef burger (RM) in red, plant-based commercial burger (PBCB) in green, plant-based home-made burger (PP) in purple, and digestion blanks group in blue. The QC samples (orange) are grouped and centered in the plot.................................................…S-24

Figure S2. PCA score plot component 1 vs component 2 for the in vitro colonic fermentation metabolic profile. Eight groups of samples were analyzed from this model: Beef burger + micro (RM) in dark red, plant-based commercial burger + micro (PBCB) in dark green, plant-based home-made burger (PP) in dark purple, their respective blanks without micro in pale red, pale green and pale purple (bl_RM, bl_PBCB, and bl_PP, respectively), blank with colon medium + micro (bl_inoculum) in dark blue, and blank with just medium (bl_medium) in pale blue. The QC samples (orange) are grouped and centered in the plot. ........................................................................S-24

Figure S3. Response of (A) phenylalanine and (B) Hydroxyproline I. Notes: RM beef meat, PBCB plant-based commercial burger, PP plant-based homemade burger, bl_inoculum, sample without any substrate. The final volume of the fermentation bottles was 70 mL. The different letters indicated the significant difference analysis for different groups at the same time point. ..........................................................................................................................................S-25

#### 1. MATERIALS AND METHODS

##### 1.1. Foodstuff material

Beef burger (125 g) was purchased from Slagerij Elings B.V. in Wageningen, consisting of 80% lean beef meat and 20% beef fat, this burger is labelled as RM. An available plant-based commercial burger, labelled as PBCB, was obtained from Albert Heijn. For the homemade plant-based burger, labelled as P.P., pea protein isolate was provided by Ingredion (Hamburg, Germany), and coconut oil was purchased from Thermo Fisher Scientific (Breda, Netherlands). The protein, fat, and water ratios were similar in the three products. More details on the composition of the foodstuff material can be bound in Table S1. The three burger types were baked in a conventional oven for 6-10 minutes at 180°C until the core reached 60°C. Before digestion, the water content of the cooked samples was measured by drying the sample using a thermal bath in sand and calculating the weight difference between the wet and dry sample.

##### 1.2. Simulated *in vitro* gastrointestinal digestion.

For the oral phase, cocked patty samples (5 g) were combined with 25 µL of $CaCl_2$ 0.3 M, 3.5 mL of SSF electrolyte stock solution, and 1.475 mL MilliQ $H_2O$ (Veolia water, Veolia Water Solutions, and Technologies Netherlands B.V.), the total volume was 10 mL. To prepare the gastric phase, the oral bolus was mixed with 7.5 mL of SGF. The pH was adjusted to 3 using 900 and 570 µL of HCl 1M for beef and plant-based analogues, respectively. Next, 5 µL of CaCl2 0.3 M were added and the volume was leveled up to 20 mL with MilliQ $H_2O$. The SGF solution was enriched with 1.6 mL of porcine pepsin stock solutions at 25 000 units $mL^{-1}$ to reach a concentration of 2000 units $mL^{-1}$ in the final chyme. Finally, the mixture was shaken for 2 h,

At the end of the gastric process, to inhibit the enzyme activity, the pH was increased to 7.0 by adding 1 mL and 450 µL of NaOH 1 M for beef and plant-based analogues respectively. Subsequently, 11 mL of SIF, 2.5 mL of fresh bile stock solution and 40 µL of $CaCl_2$ 0.3 M, 5 mL of pancreatin solution (1600 U $mL^{-1}$ amylase activity) were added, the final volume was leveled up to 40 mL by adding MilliQ H2O. At the end of the intestinal step, samples were incubated for 2 hours on a rotating device. Additionally, control samples were put through the same procedure without any digestive enzymes added. As an alternative to the enzyme solutions, MilliQ H2O was added. All intestinal digests were centrifuged to halt the enzymatic process (4 °C, 20 000 g, 10 min). Finally, 25 mL of supernatants were collected for further analysis, while pellets were freeze-dried and pooled to be used as pre-digested samples for the in vitro colonic fermentation.

#### 1.3. In-vitro batch fermentation

Colonic fermentation was carried out based on previously reported protocols, with some modifications (Huyan et al., 2022). All the information relevant to *in vitro* fermentation has been included in the Supporting Information (Section 1.3.). Briefly, fecal microbiota supernatant (FMS) was prepared by homogenizing 40.0 g of fresh feces (five donors) in 200 mL of anaerobic phosphate buffer using a stomacher bag, and a buffered colon medium consisting of different amounts of $K_2HPO_4$, $NaHCO_3$, yeast extract, peptone, mucin, L-cysteine, and Tween 80. Pre-digested samples from *in vitro* gastrointestinal digestion were mixed with buffered colon medium and FMS in 10 mL glass water-jacketed vessels. The vessels were placed in an incubator (37°C) on a rotating shaker (150 rpm). Batch cultures ran for a period of 48 hours, slurry fractions were taken at different time points (0, 6, 12, 24, and 48 hours); for donors 3, 4, and 5, slurry fractions were also collected at 3 and 32 hours. All slurry fractions were centrifuged and quenched with liquid $N_2$ immediately after sampled and stored at −20 °C until further use. A total of 8 digestions in parallel were performed per donor: the three pre-digested burgers with FMS, the three pre-digested burgers without FMS (blank samples), and two blanks of the fermentation process, with and without FMS. More details about sample composition and assigned labels can be found in **Table S2**

#### 2. TABLES AND FIGURES

*Table S1. Composition of foodstuff material*

| Burguer | Composition |
|---|---|
| Beef burger | Lean (80 %), beef fat (20%). |
| Plant-based commercial burger (PBCB) | Water, pea protein (16%), canola oil, coconut oil, rice protein, flavouring, stabilizer (methylcellulose), potato starch, apple extract, colour (beetroot red), maltodextrin, pomegranate extract, salt, potassium salt, concentrated lemon juice, maize vinegar, carrot powder, emulsifier (sunflower lecithin). |
| Homemade plant-based burger (PP) | Water, pea protein (19 %), coconut oil (19%). |

***Table S2****. Composition of the flasks subjected to the in vitro colonic fermentation. Sample labelling is in parentheses next to the treatment name.*

| Treatment names | Buffered colon medium | FMS | Pre-digested sample |
|---|---|---|---|
| Beef meat (RM) | 4.3 mL | 0.7 mL | 2 mL |
| Plant-based commercial burger (PBCB) | 4.3 mL | 0.7 mL | 2 mL |
| Plant-based homemade burger (PP) | 4.3 mL | 0.7 mL | 2 mL |
| Blank with beef meat (bl_ RM) | 4.3 mL | 0.7 mL PBS | 2 mL |
| Blank of plant-based commercial burger (bl_PCBC) | 4.3 mL | 0.7 mL PBS | 2 mL |
| Blank of plant-based homemade burger (bl_PP) | 4.3 mL | 0.7 mL PBS | 2 mL |
| Blank without micro (bl_innoculum) | 4.3 mL | 0.7 mL PBS | 2 mL Milli-Q |
| Blank with micro (bl_medium) | 4.3 mL | 0.7 mL | 2 mL Milli-Q |

*Table S3. Features of interest obtained in simulated "in vitro" gastrointestinal digestion. The feature is defined as "retention time (min)" _ "m/z". It includes the name of the compound, the formula, group, INCHIKEY, the ontology and the level of identification.*

| Feature ID | Compound name | Marker of | Formula | InChIKey | Analysis | Ontology | Level of identification |
|---|---|---|---|---|---|---|---|
| 9.462_346.0546 | Guanosine cyclic monophosphate | PBCB | C10H12N5O7P | ZOOGRGPOEVQQDX-UUOKFMHZSA-N | HILIC ESI (+) | 3',5'-cyclic purine nucleotides | Level 2a |
| 5.325_298.09671 | 5'-S-Methylthioadenosine | PBCB | C11H15N5O3S | WUUGFSXJNOTRMR-IOSLPCCCSA-N | HILIC ESI (+) | 5'-deoxy-5'-thionucleosides | Level 3b |
| 1_134.04691 | Adenine | PBCB | C5H5N5 | GFFGJBXGBJISGV-UHFFFAOYSA-N | RP ESI (-) | 6-aminopurines | Level 3b |
| 9.133_204.1241 | Acetylcarnitine | RM | C9H18NO4 | RDHQFKQIGNGIED-UHFFFAOYSA-N | HILIC ESI (+) | Acyl carnitines | Level 2a |
| 8.353_232.155 | Butyryl carnitine | RM | C11H21NO4 | LRCNOZRCYBNMEP-UHFFFAOYSA-N | HILIC ESI (+) | Acyl carnitines | Level 2a |
| 0.927_204.123 | Acetylcarnitine | RM | C9H18NO4 | RDHQFKQIGNGIED-UHFFFAOYSA-N | RP ESI (+) | Acyl carnitines | Level 2a |
| 9.346_262.12869 | O-succinylcarnitine | RM | C11H19NO6 | WEWBWVMTOYUPHH-GXUZYPEDSA-N | HILIC ESI (+) | Acyl carnitines | Level 2a (GNPS) |
| 1.261_218.13879<br>8.742_218.13901 | Propionylcarnitine | RM | C10H19NO4 | UFAHZIUFPNSHSL-UHFFFAOYSA-N | RP ESI (+)<br>HILIC ESI (+) | Acyl carnitines | Level 2a |
| 8.027_260.18549 | Hexanoyl-L-Carnitine | RM | C13H25NO4 | VVPRQWTYSNDTEA-LLVKDONJSA-N | HILIC ESI (+) | Acyl carnitines | Level 3b |
| 0.953_146.11771<br>9.409_146.11771 | Acetylcholine | RM | C7H16NO2 | OIPILFWXSMYKGL-UHFFFAOYSA-N | RP ESI (+)<br>HILIC ESI (+) | Acyl cholines | Level 3b |
| 0.952_138.054<br>9.492_138.0551 | Trigonelline | PBCB | C7H7NO2 | WWNNZCOKKKDOPX-UHFFFAOYSA-N | RP ESI (+)<br>HILIC ESI (+) | Alkaloids and derivatives | Level 3b |
| 12.21_189.1347 | Homoarginine | PBCB | C7H16N4O2 | QUOGESRFPZDMMT-YFKPBYRVSA-N | HILIC ESI (+) | Alpha amino acids | Level 2a |
| 10.011_132.0768 | Creatine | RM | C4H9N3O2 | CVSVTCORWBXHQV-UHFFFAOYSA-N | HILIC ESI (+) | Alpha amino acids and derivatives | Level 2a |

*Table S3. Features of interest obtained in simulated "in vitro" gastrointestinal digestion. The feature is defined as "retention time (min)" _ "m/z". It includes the name of the compound, the formula, group, INCHIKEY, the ontology and the level of identification.*

| Feature ID | Compound name | Marker of | Formula | InChIKey | Analysis | Ontology | Level of identification |
|---|---|---|---|---|---|---|---|
| 9.993_130.05 | 5-OXO-L-PROLINE | RM | C5H7NO3 | ODHCTXKNWHHXJC-VKHMYHEASA-N | HILIC ESI (+) | Alpha amino acids and derivatives | Level 3b |
| 9.983_90.055 | Alanine | RM | C18H36N4O4 | CSQNHSGHAPRGPQ-YTFOTSKYSA-N | HILIC ESI (+) | Amino acids | Level 4a |
| 0.912_175.11909<br>12.418_175.1194 | Arginine | PBCB | C6H14N4O2 | ODKSFYDXXFIFQN-UHFFFAOYSA-N | RP ESI (+)<br>HILIC ESI (+) | Amino acids | Level 2ª (GNPS) |
| 9.89_133.0609 | Asparagine | PBCB | C4H8N2O3 | DCXYFEDJOCDNAF-REOHCLBHSA-N | HILIC ESI (+) | Amino acids | Level 4a |
| 9.992_147.0766 | Glutamine | RM | C5H10N2O3 | ZDXPYRJPNDTMRX-VKHMYHEASA-N | HILIC ESI (+) | Amino acids | Level 2a (GNPS) |
| 1.572_130.0871<br>8.835_132.1022 | Leucine | RM | C6H13NO2 | ROHFNLRQFUQHCH-YFKPBYRVSA-N | RP ESI (-)<br>HILIC ESI (+) | Amino acids | Level 3b |
| 0.805_147.11279<br>12.45_147.1129 | Lysine | RM | C6H14N2O2 | KDXKERNSBIXSRK-UHFFFAOYSA-N | RP ESI (+)<br>HILIC ESI (+) | Amino acids | Level 2a |
| 1.159_150.0585<br>9.274_150.0584 | Methionine | RM | C5H11NO2S | FFEARJCKVFRZRR-UHFFFAOYSA-N | RP ESI (+)<br>HILIC ESI (+) | Amino acids | Level 2a |
| 2.21_164.07159<br>2.253_166.0865<br>8.659_166.0871 | Phenylalanine | RM | C9H11NO2 | COLNVLDHVKWLRT-QMMMGPOBSA-N | RP ESI (+)<br>RP ESI (-)<br>HILIC ESI (+) | Amino acids | Level 2ª (GNPS) |
| 0.969_116.0708<br>9.526_116.0708 | Proline | PBCB | C5H9NO2 | ONIBWKKTOPOVIA-UHFFFAOYSA-N | RP ESI (+)<br>HILIC ESI (+) | Amino acids | Level 3b |

*Table S3. Features of interest obtained in simulated "in vitro" gastrointestinal digestion. The feature is defined as "retention time (min)" _ "m/z". It includes the name of the compound, the formula, group, INCHIKEY, the ontology and the level of identification.*

| Feature ID | Compound name | Marker of | Formula | InChIKey | Analysis | Ontology | Level of identification |
|---|---|---|---|---|---|---|---|
| 3.491_205.0972<br>3.435_203.0827<br>8.948_205.09731 | Tryptophan | RM | C11H12N2O2 | QIVBCDIJIAJPQS-VIFPVBQESA-N | RP ESI (+)<br>RP ESI (-)<br>HILIC ESI (+) | Amino acids | Level 2ª (GNPS) |
| 1.083_118.0861 | Valine | RM | C5H11NO2 | KZSNJWFQEVHDMF-UHFFFAOYSA-N | RP ESI (+) | Amino acids | Level 2a |
| 0.969_160.13319<br>9.264_160.1335 | 4-aminovaleric acid betaine | RM | C8H17NO2 | CDLVFVFTRQPQFU-UHFFFAOYSA-N | RP ESI (+)<br>HILIC ESI (+) | Aminovaleric acid betaine | Level 2a |
| 11.741_203.15021 | ADMA | PBCB | C8H18N4O2 | YDGMGEXADBMOMJ-LURJTMIESA-N | HILIC ESI (+) | Arginine and derivatives | Level 2a |
| 15.035_112.0866 | Histamine | RM | C5H9N3 | NTYJJOPFIAHURM-UHFFFAOYSA-N | HILIC ESI (+) | Biogenic amines | Level 4a |
| 10.281_638.28729 | STL534242 | PBCB | C33H45NO10 | ACPAJGMMTDTAEW-RAAOCHHKSA-N | HILIC ESI (+) | Cardenolide glycosides and derivatives | Level 3b |
| 12.558_721.33557 | 4-((3S,10R,13R,14S,17R)-3-(((2R,3R,4S,5R,6R)-3,4-dihydroxy-6-methyl-5-(((2S,3R,4S,5S,6R)-3,4,5-trihydroxy-6-(hydroxymethyl)tetrahydro-2H-pyran-2-yl)oxy)tetrahydro-2H-pyran-2-yl)oxy)-14-hydroxy-10-(hydroxymethyl)-13-methylhexadecahydro-1H-cyclopenta[a]phenanthren-17-yl)furan-2(5H)-one | PBCB | C35H54O14 | OFSZOCPGPLMCBG-VMJITFGJSA-N | HILIC ESI (+) | Cardenolide glycosides and derivatives | Level 3b |

*Table S3. Features of interest obtained in simulated "in vitro" gastrointestinal digestion. The feature is defined as "retention time (min)" _ "m/z". It includes the name of the compound, the formula, group, INCHIKEY, the ontology and the level of identification.*

| **Feature ID** | **Compound name** | **Marker of** | **Formula** | **InChIKey** | **Analysis** | **Ontology** | **Level of identification** |
|---|---|---|---|---|---|---|---|
| 12.717_733.34668 | STL565352 | PBCB | C36H54O14 | FHIREUBIEIPPMC-YPZGHNHTSA-N | HILIC ESI (+) | Cardenolide glycosides and derivatives | Level 3b |
| 0.947_162.11259<br>10.256_162.1129 | L-Carnitine | RM | C7H15NO3 | PHIQHXFUZVPYII-ZCFIWIBFSA-N | RP ESI (+)<br>HILIC ESI (+) | Carnitines | Level 2a |
| 13.184_487.2626 | AKOS040738274 | PBCB | C24H42O7 | DQOOFQPFNNGTHL-UHFFFAOYSA-N | HILIC ESI (+) | Colensane and clerodane diterpenoids | Level 3b |
| 10.027_668.4455 | H-Val-Ile-Leu-Pro-Arg-Ala-OH | PBCB | C35H61N3O9 | OWUREPXBPJFMOK-CIRFPNLUSA-N | HILIC ESI (+) | Cyclic depsipeptides | Level 2a |
| 8.178_276.1192 | Glutamic acid-glutamine | PBCB | C10H17N3O6 | MGHKSHCBDXNTHX-UHFFFAOYSA-N | HILIC ESI (+) | Dipeptides | Level 2a |
| 1.875_322.18759<br>12.552_322.18771 | Phenylalanine-arginine | PBCB | C15H23N5O3 | OZILORBBPKKGRI-RYUDHWBXSA-N | RP ESI (+)<br>HILIC ESI (+) | Dipeptides | Level 2a (GNPS) |
| 7.721_305.0983 | Aspartate– glutamate | RM | C11H16N2O8 | OPVPGKGADVGKTG-BQBZGAKWSA-N | HILIC ESI (+) | Dipeptides | Level 2a |
| 2.884_203.1391<br>2.809_201.1244 | Alanine-(iso)leucine | RM | C9H18N2O3 | ZSOICJZJSRWNHX-ACZMJKKPSA-N | RP ESI (+)<br>RP ESI (-) | Dipeptides | Level 3b |
| 1.36_189.1234 | Alanine-valine | RM | C8H16N2O3 | LIWMQSWFLXEGMA-WDSKDSINSA-N | RP ESI (+) | Dipeptides | Level 3b |
| 1.044_288.20309 | (Iso)leucine-arginine | PBCB | C12H25N5O3 | SENJXOPIZNYLHU-UHFFFAOYSA-N | RP ESI (+) | Dipeptides | Level 2a (GNPS) |
| 8.152_287.05551 | Kaempferol | PBCB | C15H10O6 | IQPNAANSBPBGFQ-UHFFFAOYSA-N | HILIC ESI (+) | Flavones | Level 3b |
| 4.329_675.3324 | tert-butyl (1-(4-((7-acetamido-1,2,3-trimethoxy-9-oxo-5,6,7,9- | PBCB | C35H48N4O8 | IREXLBYIZXPNNP-UHFFFAOYSA-N | RP ESI (+) | Gamma amino acids and derivatives | Level 3b |

*Table S3. Features of interest obtained in simulated "in vitro" gastrointestinal digestion. The feature is defined as "retention time (min)" _ "m/z". It includes the name of the compound, the formula, group, INCHIKEY, the ontology and the level of identification.*

| Feature ID | Compound name | Marker of | Formula | InChIKey | Analysis | Ontology | Level of identification |
|---|---|---|---|---|---|---|---|
| | tetrahydrobenzo[a]heptalen-10-yl)amino)butanoyl)piperidin-4-yl)carbamate | | | | | | |
| 1.017_259.02231 | D-Glucose-6-phosphate | RM | C6H13O9P | NBSCHQHZLSJFNQ-GASJEMHNSA-N | RP ESI (-) | Hexose phosphates | Level 3b |
| 0.799_198.08749 | N,N-Acetylhistidine | RM | C8H11N3O3 | KBOJOGQFRVVWBH-ZETCQYMHSA-N | RP ESI (+) | Histidine and derivatives | Level 2a |
| 0.878_239.11481 | Anserine | RM | C10H16N4O3 | MYYIAHXIVFADCU-QMMMGPOBSA-N | RP ESI (-) | Hybrid peptides | Level 2a |
| 1.347_227.114<br>12.676_227.11459 | L-Carnosine | RM | C9H14N4O3 | CQOVPNPJLQNMDC-ZETCQYMHSA-N | RP ESI (+)<br>HILIC ESI (+) | Hybrid peptides | Level 2a |
| 0.908_407.116 | STL475082 | PBCB | C19H20N4O5 | MCCCQAKRHCMRDG-UHFFFAOYSA-N | RP ESI (+) | Hydantoins | Level 3b |
| 12.881_503.2962 | 25S-Inokosterone | PBCB | C27H44O7 | JQNVCUBPURTQPQ-BMZRUTLMSA-N | HILIC ESI (+) | Hydroxy bile acids, alcohols and derivatives | Level 3b |
| 1.291_137.0459<br>1.071_135.0314<br>7.255_137.04601 | Hypoxanthine | RM | C5H4N4O | FDGQSTZJBFJUBT-UHFFFAOYSA-N | RP ESI (+)<br>RP ESI (-)<br>HILIC ESI (+) | Hypoxanthines | Level 2a |
| 11.533_601.30737 | Ranaconitine | PBCB | C32H44N2O9 | XTSVKUJYTUPYRJ-UHFFFAOYSA-N | HILIC ESI (+) | Lappaconitine-type diterpenoid alkaloids | Level 3b |
| 14.646_670.35248 | AKOS037500753 | RM | C38H47N5O6 | OCPBZMNGFBHXLX-UHFFFAOYSA-N | HILIC ESI (+) | Leucine and derivatives | Level 3b |
| 2.505_308.18561 | O-Methacryloyl-N-octanoyl-L-tyrosine | PBCB | C17H25NO4 | OWYLAEYXIQKAOL-UHFFFAOYSA-N | HILIC ESI (+) | N-acyl amides | Level 2a |

*Table S3. Features of interest obtained in simulated "in vitro" gastrointestinal digestion. The feature is defined as "retention time (min)" _ "m/z". It includes the name of the compound, the formula, group, INCHIKEY, the ontology and the level of identification.*

| Feature ID | Compound name | Marker of | Formula | InChIKey | Analysis | Ontology | Level of identification |
|---|---|---|---|---|---|---|---|
| 8.404_301.22311 | 5-(Diaminomethylideneamino)-2-(octanoylamino)pentanoic acid | PBCB | C20H28O2 | SHGAZHPCJJPHSC-YCNIQYBTSA-N | HILIC ESI (+) | N-acyl amides | Level 2a |
| 3.819_123.0554 | Nicotinamide | RM | C6H6N2O | DFPAKSUCGFBDDF-UHFFFAOYSA-N | HILIC ESI (+) | Nicotinamides | Level 2a |
| 0.936_365.10559 | Sucrose | PBCB | C12H22O11 | CZMRCDWAGMRECN-UGDNZRGBSA-N | RP ESI (+) | O-glycosyl compounds | Level 2a |
| 2.36_373.28101 | (Iso)leucine-(iso)leucine-lysine | PBCB | C18H36N4O4 | DBEPLOCGEIEOCV-WSBQPABSSA-N | RP ESI (+) | Oligopeptides | Level 2a (GNPS) |
| 6.988_302.20761 | (Iso)leucine-Glycine-(Iso)leucine | RM | C14H27N3O4 | MQFGXJNSUJTXDT-QSFUFRPTSA-N | RP ESI (+) | Oligopeptides | Level 2a (GNPS) |
| 8.019_394.19751 | Glutamine-Valine-Phenylalanine | PBCB | C19H27N3O6 | FVGOGEGGQLNZGH-DZKIICNBSA-N | RP ESI (+) | Oligopeptides | Level 2a |
| 10.338_787.32068 | AKOS030494580 | RM | C40H46N6O11 | KRVIIMWSCXORIQ-NHKHRBQYSA-N | HILIC ESI (+) | Oligopeptides | Level 3b |
| 1.181_359.26541 | Leucine-valine-lysine | PBCB | C23H34O3 | CRRKVZVYZQXICQ-RJJCNJEVSA-N | RP ESI (+) | Oligopeptides | Level 2a |
| 9.848_522.20312 | Raffinose | PBCB | C18H32O16 | QWIZNVHXZXRPDR-WSCXOGSTSA-N | HILIC ESI (+) | Oligosaccharides | Level 2a |
| 0.943_689.21118<br>10.256_689.21143 | Stachyose | PBCB | C24H42O21 | UQZIYBXSHAGNOE-XNSRJBNMSA-N | RP ESI (+)<br>HILIC ESI (+) | Oligosaccharides | Level 2a |
| 0.907_124.0075 | Taurine | RM | C2H7NO3S | XOAAWQZATWQOTB-UHFFFAOYSA-N | RP ESI (-) | Organosulfonic acids | Level 3b |
| 12.508_487.29861 | AKOS015960725 | PBCB | C27H44O6 | UPEZCKBFRMILAV-DBDKQAFGSA-N | HILIC ESI (+) | Pentahydroxy bile acids, alcohols and derivatives | Level 2a |

*Table S3. Features of interest obtained in simulated "in vitro" gastrointestinal digestion. The feature is defined as "retention time (min)" _ "m/z". It includes the name of the compound, the formula, group, INCHIKEY, the ontology and the level of identification.*

| Feature ID | Compound name | Marker of | Formula | InChIKey | Analysis | Ontology | Level of identification |
|---|---|---|---|---|---|---|---|
| 1.231_377.08539 | melibiose | PBCB | C18H18O9 | SGQJDTLFQCDZGS-UQEKQVNISA-N | RP ESI (-) | Phenolic glycosides | Level 2a (GNPS) |
| 9.724_332.0755 | 2'-Deoxyadenosine-5'-monophosphate | PBCB | C10H14N5O6P | KHWCHTKSEGGWEX-RRKCRQDMSA-N | HILIC ESI (+) | Purine 2'-deoxyribonucleoside monophosphates | Level 3b |
| 1.312_284.0993<br>1.273_282.08469<br>8.748_284.09909 | Guanosine | PBCB | C10H13N5O5 | NYHBQMYGNKIUIF-UUOKFMHZSA-N | RP ESI (+)<br>RP ESI (-)<br>HILIC ESI (+) | Purine nucleosides | Level 2a |
| 1.291_269.08841<br>1.271_267.07349<br>8.047_269.0882 | Inosine | RM | C10H12N4O5 | UGQMRVRMYYASKQ-KQYNXXCUSA-N | RP ESI (+)<br>RP ESI (-)<br>HILIC ESI (+) | Purine nucleosides | Level 2a |
| 10.361_364.06519 | Guanosine 5'-monophosphate | PBCB | C10H14N5O8P | RQFCJASXJCIDSX-UUOKFMHZSA-N | HILIC ESI (+) | Purine ribonucleoside monophosphates | Level 2a |
| 0.999_348.07941<br>1.029_346.05679<br>9.776_348.07071 | ADENOSINE 5'-MONOPHOSPHATE | PBCB | C10H14N5O7P | UDMBCSSLTHHNCD-KQYNXXCUSA-N | RP ESI (+)<br>RP ESI (-)<br>HILIC ESI (+) | Purine ribonucleoside monophosphates | Level 3b |
| 1.291_152.0565<br>8.747_152.05679 | Guanine | PBCB | C5H5N5O | UYTPUPDQBNUYGX-UHFFFAOYSA-N | RP ESI (+)<br>HILIC ESI (+) | Purines and purine derivatives | Level 2a |
| 2.88_353.08801 | Cryptochlorogenic acid | PBCB | C16H18O9 | GYFFKZTYYAFCTR-AVXJPILUSA-N | RP ESI (-) | Quinic acids and derivatives | Level 2a |
| 1.002_322.0444<br>10.276_324.05899 | Cytidine monophosphate | PBCB | C9H14N3O8P | UOOOPKANIPLQPU-XVFCMESISA-N | RP ESI (-)<br>HILIC ESI (+) | Ribonucleoside 3'-phosphates | Level 3b |

*Table S3. Features of interest obtained in simulated "in vitro" gastrointestinal digestion. The feature is defined as "retention time (min)" _ "m/z". It includes the name of the compound, the formula, group, INCHIKEY, the ontology and the level of identification.*

| Feature ID | Compound name | Marker of | Formula | InChIKey | Analysis | Ontology | Level of identification |
|---|---|---|---|---|---|---|---|
| 8.928_761.40167 | AKOS037623077 | PBCB | C39H62O13 | GHUUTEXYFBCKSM-UYXGSHOPSA-N | HILIC ESI (+) | Steroidal saponins | Level 2a |
| 3.232_599.3512 11.198_599.35132 | Polyphyllin A | PBCB | C33H52O8 | WXMARHKAXWRNDM-GAMIEDRGSA-N | RP ESI (+) HILIC ESI (+) | Steroidal saponins | Level 3b |
| 0.893_181.0714 | D-Sorbitol | PBCB | C6H14O6 | FBPFZTCFMRRESA-NQAPHZHOSA-N | RP ESI (-) | Sugar alcohols | Level 3b |
| 1.137_191.0201 | Citric acid | PBCB | C6H8O7 | KRKNYBCHXYNGOX-UHFFFAOYSA-N | RP ESI (-) | Tricarboxylic acids and derivatives | Level 3b |
| 14.289_965.50708 | Soyasaponin Bb | PBCB | C48H78O18 | PTDAHAWQAGSZDD-IOVCITQVSA-N | RP ESI (+) | Triterpene saponins | Level 4a |
| 10.024_789.46161 | Ginsenoside Rk1 | PBCB | C42H70O12 | KWDWBAISZWOAHD-SDTJYSPYSA-N | HILIC ESI (+) | Triterpenoids | Level 2a |
| 1.71_311.1243 | gamma-Glutamyltyrosine | PBCB | C14H18N2O6 | VVLXCWVSSLFQDS-QWRGUYRKSA-N | RP ESI (+) | Tyrosine and derivatives | Level 2a |

*Table S4. Samples collected during the colonic fermentation. It includes sample name, class, time, and subject.*

| Sample name | Class | Time (hours) | Subject |
|---|---|---|---|
| Don1-RM-t0 | A_RM | 0 | Donor 1 |
| Don1-RM-t06 | A_RM | 6 | Donor 1 |
| Don1-RM-t12 | A_RM | 12 | Donor 1 |
| Don1-RM-t24 | A_RM | 24 | Donor 1 |
| Don1-RM-t48 | A_RM | 48 | Donor 1 |
| Don2-RM-t0 | A_RM | 0 | Donor 2 |
| Don2-RM-t06 | A_RM | 6 | Donor 2 |
| Don2-RM-t12 | A_RM | 12 | Donor 2 |
| Don2-RM-t24 | A_RM | 24 | Donor 2 |
| Don2-RM-t48 | A_RM | 48 | Donor 2 |
| Don3-RM-t0 | A_RM | 0 | Donor 3 |
| Don3-RM-t03 | A_RM | 3 | Donor 3 |
| Don3-RM-t06 | A_RM | 6 | Donor 3 |
| Don3-RM-t12 | A_RM | 12 | Donor 3 |
| Don3-RM-t24 | A_RM | 24 | Donor 3 |
| Don3-RM-t32 | A_RM | 32 | Donor 3 |
| Don3-RM-t48 | A_RM | 48 | Donor 3 |
| Don4-RM-t0 | A_RM | 0 | Donor 4 |
| Don4-RM-t03 | A_RM | 3 | Donor 4 |
| Don4-RM-t06 | A_RM | 6 | Donor 4 |
| Don4-RM-t12 | A_RM | 12 | Donor 4 |
| Don4-RM-t24 | A_RM | 24 | Donor 4 |
| Don4-RM-t36 | A_RM | 32 | Donor 4 |
| Don4-RM-t48 | A_RM | 48 | Donor 4 |
| Don5-RM-t0 | A_RM | 0 | Donor 5 |
| Don5-RM-t03 | A_RM | 3 | Donor 5 |
| Don5-RM-t06 | A_RM | 6 | Donor 5 |
| Don5-RM-t12 | A_RM | 12 | Donor 5 |
| Don5-RM-t24 | A_RM | 24 | Donor 5 |
| Don5-RM-t31 | A_RM | 32 | Donor 5 |
| Don5-RM-t48 | A_RM | 48 | Donor 5 |
| Don1-bl_RM-t0 | B_bl_RM | 0 | Donor 1 |
| Don1-bl_RM-t06 | B_bl_RM | 6 | Donor 1 |
| Don1-bl_RM-t12 | B_bl_RM | 12 | Donor 1 |
| Don1-bl_RM-t24 | B_bl_RM | 24 | Donor 1 |
| Don1-bl_RM-t48 | B_bl_RM | 48 | Donor 1 |
| Don2-bl_RM-t0 | B_bl_RM | 0 | Donor 2 |
| Don2-bl_RM-t06 | B_bl_RM | 6 | Donor 2 |

*Table S4. Samples collected during the colonic fermentation. It includes sample name, class, time, and subject.*

| Sample name | Class | Time (hours) | Subject |
|---|---|---|---|
| Don2-bl_RM-t12 | B_bl_RM | 12 | Donor 2 |
| Don2-bl_RM-t24 | B_bl_RM | 24 | Donor 2 |
| Don2-bl_RM-t48 | B_bl_RM | 48 | Donor 2 |
| Don3-bl_RM-t0 | B_bl_RM | 0 | Donor 3 |
| Don3-bl_RM-t03 | B_bl_RM | 3 | Donor 3 |
| Don3-bl_RM-t06 | B_bl_RM | 6 | Donor 3 |
| Don3-bl_RM-t12 | B_bl_RM | 12 | Donor 3 |
| Don3-bl_RM-t24 | B_bl_RM | 24 | Donor 3 |
| Don4-bl_RM-t0 | B_bl_RM | 0 | Donor 4 |
| Don4-bl_RM-t03 | B_bl_RM | 3 | Donor 4 |
| Don4-bl_RM-t06 | B_bl_RM | 6 | Donor 4 |
| Don4-bl_RM-t12 | B_bl_RM | 12 | Donor 4 |
| Don4-bl_RM-t24 | B_bl_RM | 24 | Donor 4 |
| Don4-bl_RM-t36 | B_bl_RM | 32 | Donor 4 |
| Don4-bl_RM-t48 | B_bl_RM | 48 | Donor 4 |
| Don5-bl_RM-t0 | B_bl_RM | 0 | Donor 5 |
| Don5-bl_RM-t06 | B_bl_RM | 6 | Donor 5 |
| Don5-bl_RM-t12 | B_bl_RM | 12 | Donor 5 |
| Don5-bl_RM-t24 | B_bl_RM | 24 | Donor 5 |
| Don5-bl_RM-t31 | B_bl_RM | 32 | Donor 5 |
| Don5-bl_RM-t48 | B_bl_RM | 48 | Donor 5 |
| Don1-Bey-t0 | C_PBCB | 0 | Donor 1 |
| Don1-Bey-t06 | C_PBCB | 6 | Donor 1 |
| Don1-Bey-t12 | C_PBCB | 12 | Donor 1 |
| Don1-Bey-t24 | C_PBCB | 24 | Donor 1 |
| Don1-Bey-t48 | C_PBCB | 48 | Donor 1 |
| Don2-Bey-t0 | C_PBCB | 0 | Donor 2 |
| Don2-Bey-t06 | C_PBCB | 6 | Donor 2 |
| Don2-Bey-t12 | C_PBCB | 12 | Donor 2 |
| Don2-Bey-t24 | C_PBCB | 24 | Donor 2 |
| Don2-Bey-t48 | C_PBCB | 48 | Donor 2 |
| Don3-Bey-t0 | C_PBCB | 0 | Donor 3 |
| Don3-Bey-t03 | C_PBCB | 3 | Donor 3 |
| Don3-Bey-t06 | C_PBCB | 6 | Donor 3 |
| Don3-Bey-t12 | C_PBCB | 12 | Donor 3 |
| Don3-Bey-t24 | C_PBCB | 24 | Donor 3 |
| Don3-Bey-t32 | C_PBCB | 32 | Donor 3 |
| Don3-Bey-t48 | C_PBCB | 48 | Donor 3 |

*Table S4. Samples collected during the colonic fermentation. It includes sample name, class, time, and subject.*

| Sample name | Class | Time (hours) | Subject |
|---|---|---|---|
| Don4-Bey-t0 | C_PBCB | 0 | Donor 4 |
| Don4-Bey-t03 | C_PBCB | 3 | Donor 4 |
| Don4-Bey-t06 | C_PBCB | 6 | Donor 4 |
| Don4-Bey-t12 | C_PBCB | 12 | Donor 4 |
| Don4-Bey-t24 | C_PBCB | 24 | Donor 4 |
| Don4-Bey-t36 | C_PBCB | 32 | Donor 4 |
| Don4-Bey-t48 | C_PBCB | 48 | Donor 4 |
| Don5-Bey-t0 | C_PBCB | 0 | Donor 5 |
| Don5-Bey-t03 | C_PBCB | 3 | Donor 5 |
| Don5-Bey-t06 | C_PBCB | 6 | Donor 5 |
| Don5-Bey-t12 | C_PBCB | 12 | Donor 5 |
| Don5-Bey-t24 | C_PBCB | 24 | Donor 5 |
| Don5-Bey-t31 | C_PBCB | 32 | Donor 5 |
| Don5-Bey-t48 | C_PBCB | 48 | Donor 5 |
| Don1-bl_Bey-t0 | D_bl_PBCB | 0 | Donor 1 |
| Don1-bl_Bey-t06 | D_bl_PBCB | 6 | Donor 1 |
| Don1-bl_Bey-t12 | D_bl_PBCB | 12 | Donor 1 |
| Don1-bl_Bey-t24 | D_bl_PBCB | 24 | Donor 1 |
| Don1-bl_Bey-t48 | D_bl_PBCB | 48 | Donor 1 |
| Don2-bl_Bey-t0 | D_bl_PBCB | 0 | Donor 2 |
| Don2-bl_Bey-t06 | D_bl_PBCB | 6 | Donor 2 |
| Don2-bl_Bey-t12 | D_bl_PBCB | 12 | Donor 2 |
| Don2-bl_Bey-t24 | D_bl_PBCB | 24 | Donor 2 |
| Don2-bl_Bey-t48 | D_bl_PBCB | 48 | Donor 2 |
| Don3-bl_Bey-t0 | D_bl_PBCB | 0 | Donor 3 |
| Don3-bl_Bey-t03 | D_bl_PBCB | 3 | Donor 3 |
| Don3-bl_Bey-t06 | D_bl_PBCB | 6 | Donor 3 |
| Don3-bl_Bey-t12 | D_bl_PBCB | 12 | Donor 3 |
| Don3-bl_Bey-t24 | D_bl_PBCB | 24 | Donor 3 |
| Don4-bl_Bey-t0 | D_bl_PBCB | 0 | Donor 4 |
| Don4-bl_Bey-t03 | D_bl_PBCB | 3 | Donor 4 |
| Don4-bl_Bey-t06 | D_bl_PBCB | 6 | Donor 4 |
| Don4-bl_Bey-t12 | D_bl_PBCB | 12 | Donor 4 |
| Don4-bl_Bey-t24 | D_bl_PBCB | 24 | Donor 4 |
| Don4-bl_Bey-t36 | D_bl_PBCB | 32 | Donor 4 |
| Don4-bl_Bey-t48 | D_bl_PBCB | 48 | Donor 4 |
| Don5-bl_Bey-t0 | D_bl_PBCB | 0 | Donor 5 |
| Don5-bl_Bey-t03 | D_bl_PBCB | 3 | Donor 5 |

*Table S4. Samples collected during the colonic fermentation. It includes sample name, class, time, and subject.*

| Sample name | Class | Time (hours) | Subject |
|---|---|---|---|
| Don5-bl_Bey-t06 | D_bl_PBCB | 6 | Donor 5 |
| Don5-bl_Bey-t12 | D_bl_PBCB | 12 | Donor 5 |
| Don5-bl_Bey-t24 | D_bl_PBCB | 24 | Donor 5 |
| Don5-bl_Bey-t31 | D_bl_PBCB | 32 | Donor 5 |
| Don5-bl_Bey-t48 | D_bl_PBCB | 48 | Donor 5 |
| Don1-PP-t0 | E_PP | 0 | Donor 1 |
| Don1-PP-t06 | E_PP | 6 | Donor 1 |
| Don1-PP-t12 | E_PP | 12 | Donor 1 |
| Don1-PP-t24 | E_PP | 24 | Donor 1 |
| Don1-PP-t48 | E_PP | 48 | Donor 1 |
| Don2-PP-t0 | E_PP | 0 | Donor 2 |
| Don2-PP-t06 | E_PP | 6 | Donor 2 |
| Don2-PP-t12 | E_PP | 12 | Donor 2 |
| Don2-PP-t24 | E_PP | 24 | Donor 2 |
| Don2-PP-t48 | E_PP | 48 | Donor 2 |
| Don3-PP-t0 | E_PP | 0 | Donor 3 |
| Don3-PP-t03 | E_PP | 3 | Donor 3 |
| Don3-PP-t06 | E_PP | 6 | Donor 3 |
| Don3-PP-t12 | E_PP | 12 | Donor 3 |
| Don3-PP-t24 | E_PP | 24 | Donor 3 |
| Don3-PP-t32 | E_PP | 32 | Donor 3 |
| Don3-PP-t48 | E_PP | 48 | Donor 3 |
| Don4-PP-t0 | E_PP | 0 | Donor 4 |
| Don4-PP-t03 | E_PP | 3 | Donor 4 |
| Don4-PP-t06 | E_PP | 6 | Donor 4 |
| Don4-PP-t12 | E_PP | 12 | Donor 4 |
| Don4-PP-t24 | E_PP | 24 | Donor 4 |
| Don4-PP-t36 | E_PP | 32 | Donor 4 |
| Don4-PP-t48 | E_PP | 48 | Donor 4 |
| Don5-PP-t0 | E_PP | 0 | Donor 5 |
| Don5-PP-t03 | E_PP | 3 | Donor 5 |
| Don5-PP-t06 | E_PP | 6 | Donor 5 |
| Don5-PP-t12 | E_PP | 12 | Donor 5 |
| Don5-PP-t24 | E_PP | 24 | Donor 5 |
| Don5-PP-t31 | E_PP | 32 | Donor 5 |
| Don5-PP-t48 | E_PP | 48 | Donor 5 |
| Don1-bl_PP-t0 | F_bl_PP | 0 | Donor 1 |
| Don1-bl_PP-t06 | F_bl_PP | 6 | Donor 1 |

*Table S4. Samples collected during the colonic fermentation. It includes sample name, class, time, and subject.*

| Sample name | Class | Time (hours) | Subject |
|---|---|---|---|
| Don1-bl_PP-t12 | F_bl_PP | 12 | Donor 1 |
| Don1-bl_PP-t24 | F_bl_PP | 24 | Donor 1 |
| Don1-bl_PP-t48 | F_bl_PP | 48 | Donor 1 |
| Don2-bl_PP-t0 | F_bl_PP | 0 | Donor 2 |
| Don2-bl_PP-t06 | F_bl_PP | 6 | Donor 2 |
| Don2-bl_PP-t12 | F_bl_PP | 12 | Donor 2 |
| Don2-bl_PP-t24 | F_bl_PP | 24 | Donor 2 |
| Don2-bl_PP-t48 | F_bl_PP | 48 | Donor 2 |
| Don3-bl_PP-t0 | F_bl_PP | 0 | Donor 3 |
| Don3-bl_PP-t03 | F_bl_PP | 3 | Donor 3 |
| Don3-bl_PP-t06 | F_bl_PP | 6 | Donor 3 |
| Don3-bl_PP-t12 | F_bl_PP | 12 | Donor 3 |
| Don4-bl_PP-t0 | F_bl_PP | 0 | Donor 4 |
| Don4-bl_PP-t03 | F_bl_PP | 3 | Donor 4 |
| Don4-bl_PP-t06 | F_bl_PP | 6 | Donor 4 |
| Don4-bl_PP-t12 | F_bl_PP | 12 | Donor 4 |
| Don4-bl_PP-t24 | F_bl_PP | 24 | Donor 4 |
| Don4-bl_PP-t36 | F_bl_PP | 32 | Donor 4 |
| Don4-bl_PP-t48 | F_bl_PP | 48 | Donor 4 |
| Don5-bl_PP-t0 | F_bl_PP | 0 | Donor 5 |
| Don5-bl_PP-t03 | F_bl_PP | 3 | Donor 5 |
| Don5-bl_PP-t06 | F_bl_PP | 6 | Donor 5 |
| Don5-bl_PP-t12 | F_bl_PP | 12 | Donor 5 |
| Don5-bl_PP-t24 | F_bl_PP | 24 | Donor 5 |
| Don5-bl_PP-t31 | F_bl_PP | 32 | Donor 5 |
| Don5-bl_PP-t48 | F_bl_PP | 48 | Donor 5 |
| Don1-bl_innoc-t0 | G_bl_innoc | 0 | Donor 1 |
| Don1-bl_innoc-t06 | G_bl_innoc | 6 | Donor 1 |
| Don1-bl_innoc-t12 | G_bl_innoc | 12 | Donor 1 |
| Don1-bl_innoc-t24 | G_bl_innoc | 24 | Donor 1 |
| Don1-bl_innoc-t48 | G_bl_innoc | 48 | Donor 1 |
| Don2-bl_innoc-t0 | G_bl_innoc | 0 | Donor 2 |
| Don2-bl_innoc-t06 | G_bl_innoc | 6 | Donor 2 |
| Don2-bl_innoc-t12 | G_bl_innoc | 12 | Donor 2 |
| Don2-bl_innoc-t24 | G_bl_innoc | 24 | Donor 2 |
| Don2-bl_innoc-t48 | G_bl_innoc | 48 | Donor 2 |
| Don3-bl_innoc-t0 | G_bl_innoc | 0 | Donor 3 |
| Don3-bl_innoc-t03 | G_bl_innoc | 3 | Donor 3 |

*Table S4. Samples collected during the colonic fermentation. It includes sample name, class, time, and subject.*

| Sample name | Class | Time (hours) | Subject |
|---|---|---|---|
| Don4-bl_innoc-t0 | G_bl_innoc | 0 | Donor 4 |
| Don4-bl_innoc-t03 | G_bl_innoc | 3 | Donor 4 |
| Don4-bl_innoc-t06 | G_bl_innoc | 6 | Donor 4 |
| Don4-bl_innoc-t12 | G_bl_innoc | 12 | Donor 4 |
| Don4-bl_innoc-t24 | G_bl_innoc | 24 | Donor 4 |
| Don4-bl_innoc-t36 | G_bl_innoc | 32 | Donor 4 |
| Don4-bl_innoc-t48 | G_bl_innoc | 48 | Donor 4 |
| Don5-bl_innoc-t0 | G_bl_innoc | 0 | Donor 5 |
| Don5-bl_innoc-t03 | G_bl_innoc | 3 | Donor 5 |
| Don5-bl_innoc-t06 | G_bl_innoc | 6 | Donor 5 |
| Don5-bl_innoc-t12 | G_bl_innoc | 12 | Donor 5 |
| Don5-bl_innoc-t24 | G_bl_innoc | 24 | Donor 5 |
| Don5-bl_innoc-t31 | G_bl_innoc | 32 | Donor 5 |
| Don5-bl_innoc-t48 | G_bl_innoc | 48 | Donor 5 |
| Don1-bl_medium-t0 | H_bl_medium | 0 | Donor 1 |
| Don1-bl_medium-t06 | H_bl_medium | 6 | Donor 1 |
| Don1-bl_medium-t12 | H_bl_medium | 12 | Donor 1 |
| Don1-bl_medium-t24 | H_bl_medium | 24 | Donor 1 |
| Don1-bl_medium-t48 | H_bl_medium | 48 | Donor 1 |
| Don2-bl_medium-t0 | H_bl_medium | 0 | Donor 2 |
| Don2-bl_medium-t06 | H_bl_medium | 6 | Donor 2 |
| Don2-bl_medium-t12 | H_bl_medium | 12 | Donor 2 |
| Don2-bl_medium-t24 | H_bl_medium | 24 | Donor 2 |
| Don2-bl_medium-t48 | H_bl_medium | 48 | Donor 2 |
| Don3-bl_medium-t0 | H_bl_medium | 0 | Donor 3 |
| Don3-bl_medium-t03 | H_bl_medium | 3 | Donor 3 |
| Don3-bl_medium-t06 | H_bl_medium | 6 | Donor 3 |
| Don3-bl_medium-t12 | H_bl_medium | 12 | Donor 3 |
| Don3-bl_medium-t24 | H_bl_medium | 24 | Donor 3 |
| Don3-bl_medium-t48 | H_bl_medium | 48 | Donor 3 |
| Don4-bl_medium-t0 | H_bl_medium | 0 | Donor 4 |
| Don4-bl_medium-t03 | H_bl_medium | 3 | Donor 4 |
| Don4-bl_medium-t06 | H_bl_medium | 6 | Donor 4 |
| Don4-bl_medium-t12 | H_bl_medium | 12 | Donor 4 |
| Don4-bl_medium-t24 | H_bl_medium | 24 | Donor 4 |
| Don4-bl_medium-t48 | H_bl_medium | 48 | Donor 4 |
| Don5-bl_medium-t0 | H_bl_medium | 0 | Donor 5 |
| Don5-bl_medium-t03 | H_bl_medium | 3 | Donor 5 |

*Table S4. Samples collected during the colonic fermentation. It includes sample name, class, time, and subject.*

| Sample name | Class | Time (hours) | Subject |
|---|---|---|---|
| Don5-bl_medium-t12 | H_bl_medium | 12 | Donor 5 |
| Don5-bl_medium-t24 | H_bl_medium | 24 | Donor 5 |
| Don5-bl_medium-t31 | H_bl_medium | 32 | Donor 5 |
| Don5-bl_medium-t48 | H_bl_medium | 48 | Donor 5 |

*Table S5. Features of interest obtained in colonic fermentation. The feature is defined as "retention time (min)" _ "m/z". It includes the name of the compound, the formula, the INCHIKEY, the ontology and the level of identification.*

| Feature ID | Compound name | Formula | InChIKey | Analysis | Ontology | Level of identification |
|---|---|---|---|---|---|---|
| 2.233_164.07159<br>2.259_166.0863 | Phenylalanine | C9H11NO2 | COLNVLDHVKWLRT-QMMMGPOBSA-N | RP ESI (-)<br>RP ESI(+) | Amino acids | Level 2a |
| 9.408_182.082 | Tyrosine | C9H11NO3 | ZWEHNKRNPOVVGH-UHFFFAOYSA-N | HILIC ESI (+) | Amino acids | Level 2a |
| 1.156_150.0585<br>9.048_150.0587 | Methionine | C5H11NO2S | FFEARJCKVFRZRR-UHFFFAOYSA-N | RP ESI (+)<br>HILIC ESI(+) | Amino acids | Level 2a |
| 3.535_188.071<br>3.535_205.09731 | L-Tryptophan | C9H11NO2 | QIVBCDIJIAJPQS-UHFFFAOYSA-N | RP ESI (+) | Amino acids | Level 2a (GNPS) |
| 12.277_170.0923 | Methyl-Histidine | C7H11N3O2 | | HILIC ESI (+) | Histidine derivates | Level 2a |
| 12.831_189.1597 | N,N,N-Trimethyllysine | C9H20N2O2 | MXNRLFUSFKVQSK-QMMMGPOBSA-N | HILIC ESI (+) | L-alpha-amino acids | Level 2a |
| 3.276_132.06551 | Hydroxyproline I | C5H9NO3 | | RP ESI (+) | Proline derivates | Level 3b |
| 9.97_132.07671 | Hydroxyproline II | C5H9NO3 | | HILIC ESI (+) | Proline derivates | Level 2a |
| 2.384_292.1907 | C8:0-phenylalanine | C17H25NO3 | XNWZOFXSPQXPRE-HNNXBMFYSA-N | HILIC ESI (+) | N-acyl amides | Level 2a (GNPS) |
| 2.321_428.31619 | C18:2-phenylalanine | C27H45NO5 | HKUGGWURERXWKM-ABDCMALPSA-N | HILIC ESI (+) | N-acyl amides | Level 2a (GNPS) |
| 2.358_426.29999 | C18:3-pheynlalanine | C27H39NO3 | IJUCNQRDKBWNBS-GMOGWPNXSA-N | HILIC ESI (+) | N-acyl amides | Level 2a (GNPS) |
| 2.292_394.3317 | C18:2-leucine | C24H40O3 | CICRQSYFFGMUEH-FQPCFBMWSA-N | HILIC ESI (+) | N-acyl amides | Level 2a (GNPS) |
| 1.031_204.12309<br>9.09_204.1234 | Acetylcarnitine | C9H18NO4 | RDHQFKQIGNGIED-UHFFFAOYSA-N | RP ESI (+)<br>HILIC ESI(+) | Acyl carnitines | Level 2a |
| 2.913_201.12421<br>2.962_203.13921 | Alanine-(iso)leucine | C9H18N2O3 | | RP ESI (-)<br>RP ESI(+) | Dipeptides | Level 2a |

*Table S5. Features of interest obtained in colonic fermentation. The feature is defined as "retention time (min)" _ "m/z". It includes the name of the compound, the formula, the INCHIKEY, the ontology and the level of identification.*

| Feature ID | Compound name | Formula | InChIKey | Analysis | Ontology | Level of identificatio n |
|---|---|---|---|---|---|---|
| 1.482_288.20331<br>12.934_288.2034 | Arginine-(iso)leucine | C16H21NO2 | | RP ESI (+)<br>HILIC ESI(+) | Dipeptides | Level 2a (GNPS) |
| 1.313_269.16119<br>11.127_269.16071 | Histidine-(iso)leucine | C15H22N2O | | RP ESI (+)<br>HILIC ESI(+) | Dipeptides | Level 2a (GNPS) |
| 3.86_263.1427 | (Iso)leucine-methionine | C19H20O2 | | RP ESI (+) | Dipeptides | Level 2a (GNPS) |
| 1.419_260.1972<br>12.837_260.1969 | Lysine-(Iso)leucine | C13H26NO4 | | RP ESI (+)<br>HILIC ESI(+) | Dipeptides | Level 2a (GNPS) |
| 5.539_241.11909 | Pyroglutamyl-(iso)leucine | C11H18N2O4 | | RP ESI (-) | Dipeptides | Level 2a |
| 1.939_322.18771<br>12.634_322.1872 9 | Phenylalanine-arginine | C20H23N3O | | RP ESI (+)<br>HILIC ESI(+) | Dipeptides | Level 2a |
| 12.737_241.1293 | L-Anserine | C10H16N4O3 | MYYIAHXIVFADCU-QMMMGPOBSA-N | HILIC ESI (+) | Hybrid peptides | Level 2a |
| 0.852_225.0992<br>12.754_227.1138 | L-Carnosine | C9H14N4O3 | CQOVPNPJLQNMDC-ZETCQYMHSA-N | RP ESI (-)<br>HILIC ESI(+) | Hybrid peptides | Level 2a |
| 3.069_259.1297<br>3.112_261.14471 | Gamma-glutamyl-leucine | C11H20N2O5 | MYFMARDICOWMQP-YUMQZZPRSA-N | RP ESI (-)<br>HILIC ESI(+) | Dipeptides | Level 2a |
| 3.07_305.08109 | Gamma-glutamyl-S-(1-propenyl)cysteine sulfoxide | C11H18N2O6S | LMNDKWXDMBGGAL-UHFFFAOYSA-N | RP ESI (-) | Dipeptides | Level 3b |
| 1.187_277.0863 | gamma-Glutamylmethionine | C10H18N2O5 S | RQNSKRXMANOPQY-BQBZGAKWSA-N | RP ESI (-) | Dipeptides | Level 3b |

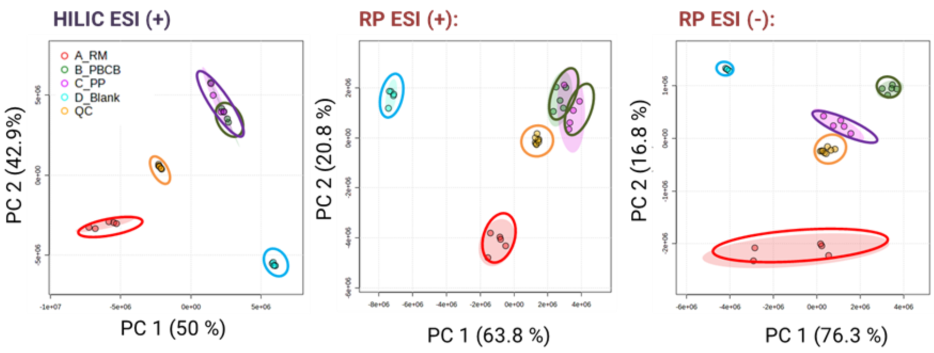

**Figure S1.** *PCA score plot component 1 vs component 2 for the simulated gastrointestinal digestion metabolic profiles, mirroring the small intestine content. Four groups of samples were analyzed from this in vitro digestion model: Beef burger (RM) in red, plant-based commercial burger (PBCB) in green, plant-based home-made burger (PP) in purple, and digestion blanks group in blue. The QC samples (orange) are grouped and centered in the plot.*

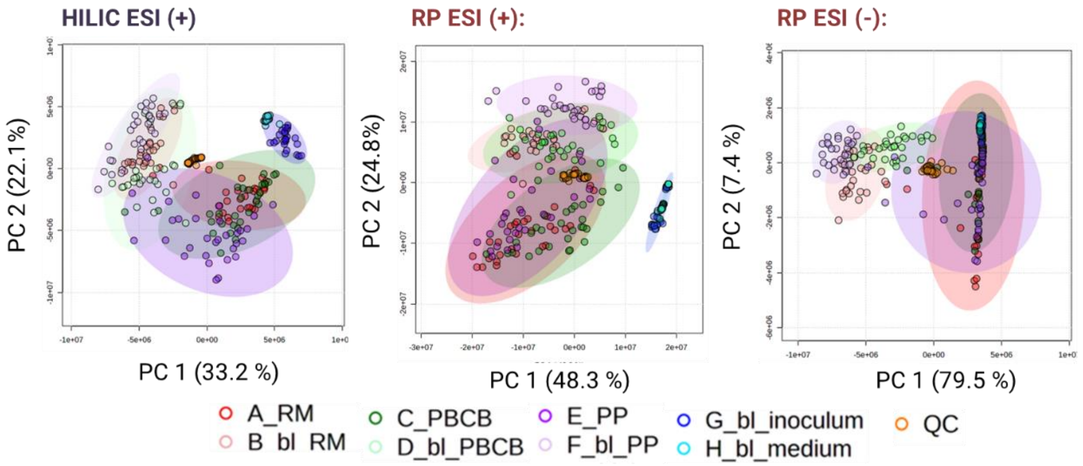

**Figure S2.** *PCA score plot component 1 vs component 2 for the in vitro colonic fermentation metabolic profile. Eight groups of samples were analyzed from this model: Beef burger + micro (RM) in dark red, plant-based commercial burger + micro (PBCB) in dark green, plant-based home-made burger (PP) in dark purple, their respective blanks without micro in pale red, pale green and pale purple (bl_RM, bl_PBCB, and bl_PP, respectively), blank with colon medium + micro (bl_inoculum) in dark blue, and blank with just medium (bl_medium) in pale blue. The QC samples (orange) are grouped and centered in the plot.*

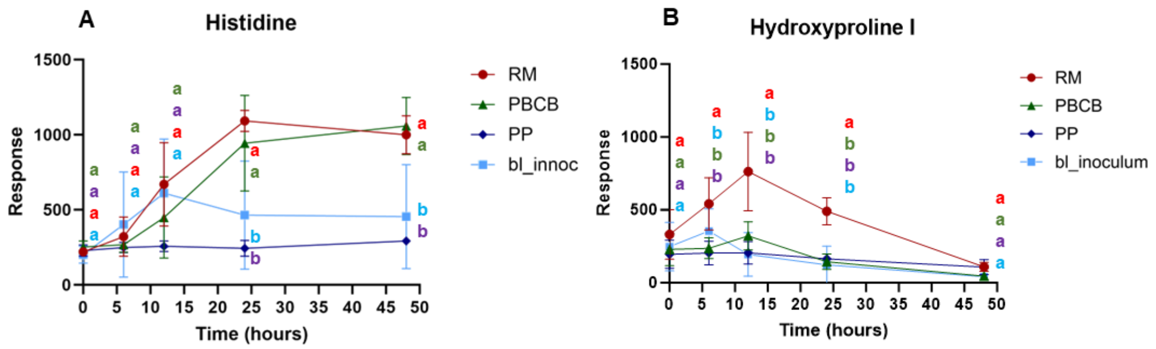

**Figure S3.** *Response of (A) phenylalanine and (B) Hydroxyproline I. Notes: RM beef meat, PBCB plant-based commercial burger, PP plant-based homemade burger, bl_inoculum, sample without any substrate. The final volume of the fermentation bottles was 70 mL. The different letters indicated the significant difference analysis for different groups at the same time point.*
